# Homodimerized Arpins: Binding to Arp2/3 Complexes with Positive Synergy to Inhibit Lamellipodia-Dependent Tumor Metastasis

**DOI:** 10.1101/2025.06.24.661301

**Authors:** Liru Liu, Xuejiao Zhang, Yun Zhu, Mengchen Pu, Yuru Geng, Min Wang, Rongguang Zhang, Shen Ge, Sheng Ye

## Abstract

The Arp2/3 complex is a central regulator of actin polymerization dynamics and cancer cell motility, with its dysregulation implicated in metastasis and therapeutic resistance. Despite its therapeutic potential, targeting Arp2/3 remains challenging due to the lack of precise molecular insights into endogenous inhibitors. Arpin, a lamellipodia-localized Arp2/3 inhibitor, has emerged as a critical modulator of cancer migration; however, the structural and mechanistic basis of its inhibitory function remains unresolved. Here, we report the 1.65-Å crystal structure of Arpin’s N-terminal globular domain, revealing a unique structural fold distinct from all known Arp2/3-binding factors. Structural and biophysical analyses demonstrate that Arpin forms homodimers via a conserved interface, which is essential for cooperative inhibition of Arp2/3-dependent actin polymerization. Functional studies in cancer models reveal that Arpin homodimerization potently suppresses lamellipodia-driven migration and invasion, effects further enhanced by engineered dimeric constructs or artificially designed dual-tailed peptides. These findings elucidate a structural basis for Arpin-mediated suppression of cancer cell motility and identify homodimeric cooperativity as a druggable mechanism for targeting metastatic progression. Our work provides a molecular framework for designing Arpin-inspired inhibitors to counteract therapeutic resistance and metastasis in cancers reliant on Arp2/3-driven actin dynamics.

## Introduction

Malignant tumor cells exploit their intrinsic migratory capabilities to invade adjacent tissues, ultimately leading to metastasize^1^. Cell migration is a sequential and interrelated multi-step process that requires the dynamic reorganization of the actin cytoskeleton, wherein the microfilaments provide mechanical support for anterior protrusion and posterior retraction^2,3^. As the rate-limiting step in actin polymerization, spontaneous nucleation constitutes a critical node in surmounting the kinetic barrier to filament initiation^4,5^. The actin-related protein (Arp) 2/3 complex remains the sole identified modulator of actin branching, which binds to the lateral surface or terminal region pre-existing actin filaments to instigate the formation of nascent daughter filaments emanating from parental filaments^6–8^. The activators of Arp2/3 complex are known as nucleation-promoting factors (NPFs), the majority of which belong to the WASP/WAVE family and share a characteristic VCA domain^9^. To tightly regulate its potent nucleation effect, the Arp2/3 complex is usually properly inhibited at distinct subcellular locations by multiple binding partners, including Coronin, Glia Maturation Factor (GMF) and Gadkin. Recently, Arpin, an Arp2/3 complex inhibitor localized to lamellipodia, was identified in a bioinformatic search for the so-called “A” motif of VCA domain and shown to suppress cancer cell migration^10^. Notably, Arpin’s expression in tumor cells, particularly breast cancer cells, is significantly downregulated compared to normal tissues^11–14^. However, the inhibitory function of Arpin appears to be somewhat controversial: initial studies showed that, instead of preventing lamellipodia protrusion, Arpin merely reduces cell migration speed and promotes cell steering^10^; while a subsequent study demonstrated that Arpin knockout cells exhibited no obvious defect in chemotaxis, indicating a weak association with cell steering^15^. The complicated and conflicting findings indicate that the precise inhibitory mechanism of Arpin remains poorly elucidated.

Arpin and NPFs share the conserved acidic (“A”) motif, which, together with a verprolin homology motif (“V”) and a central motif (“C”), compose the VCA domain. In their unbound state, all three motifs adopt disordered conformations, but they form ordered secondary structures when bound to actin or the Arp2/3 complex. Among these motifs, “V” and “C” are capable of binding to actin, while “C” and “A” can combine Arp2/3 complex at certain sites, consequently bridging the Arp2/3 complex and the first actin subunit of a daughter filament^16–18^. Arpin, which possesses a “A” motif, its C-terminal acidic tail (AT), but lacks the “V” or “C” motif, has been proved to be a competitive inhibitor of the Arp2/3 complex against VCA-domain-containing NPFs^10^. Truncation of the C-terminal AT significantly depletes the binding affinity between Arpin and Arp2/3, whereas the AT segment alone is sufficient to mediate Arpin’s inhibitory activity^10,19^. The functional findings led to an intriguing paradox: the N-terminal region of Arpin, although lacking functional contribution to Arp2/3 binding, exhibits striking sequence conservation across species. This evolutionary conservation of the N-terminal region, juxtaposed against its functional redundancy in Arp2/3 interactions, underscores the limitations imposed by the paucity of high-resolution structural data.

To fully understand its inhibitory mechanism, Arpin has been subjected to structural studies with diverse methodologies. Despite sharing a functional AT region, Arpin exhibits no discernible sequence homology to canonical NPFs such as WASP, WAVE, or WHAMM, of which the structural arrangement have been extensively characterized in complex with the Arp2/3 complex^8,20,24,25^. Furthermore, comprehensive sequence alignment against the Protein Data Bank (PDB) revealed no detectable sequence similarities, indicating that Arpin might adopt a novel fold. Nevertheless, prior to this study, no detailed structural data on intact Arpin were available. Only a low-resolution small-angle X-ray scattering (SAXS) structure has been reported, depicting a tadpole-like contour of full-length Arpin^19^. In the same study, the authors solved the complex structure of Arpin’s AT in complex with the ankyrin repeats domain of Tankyrase 2 (PDB entry: 4Z68), revealing a disordered conformation of the AT. Based on these findings, Arpin is considered to comprise two portions: an ellipsoid-shaped N-terminal globular domain and an extended linear C-terminal tail (CT).

In addition to characterizing Arpin’s full-length structure, structurally deciphering its complex with Arp2/3 has also been a focal point for researchers aiming to elucidate the molecular mechanisms underlying their interactions. Given that both Arpin and NPF utilize a conserved acidic motif to associate with the Arp2/3 complex, it is plausible that they share overlapping AT-binding sites. The cryo-electron microscopy (cryo-EM) structures of the NPF: Arp2/3 complex (PDB entries: 6UHC, 9DLX) clearly unveiled two distinct AT-binding sites: one localized on the Arp3 subunit (designated ‘site-A3’ in this work) and the other within the cleft formed by subunits Arp2 and ArpC1 (designated ‘site-A2C1’ in this work)^20,21^. Consistently, a low-resolution cryo-EM structure of the Arpin-bound Arp2/3 complex suggested the potential existence of multiple Arpin-binding sites^22^. Nevertheless, in 2022, Fregoso et al.^23^ reported the cryo-EM structure of the Arp2/3 complex with only one bound Arpin’s CT at site-A3, whereas no associated density was observed at site-A2C1 (PDB entry: 7JPN). This observation was corroborated by isothermal titration calorimetry (ITC) data from the same study, which revealed that both full-length Arpin and its CT associated with the Arp2/3 complex in a 1:1 stoichiometric ratio^23^. These conflicting findings highlight the inherent complexity of the binding mode between Arpin and the Arp2/3 complex. If both site-A3 and site-A2C1 are indeed utilized by Arpin, their binding affinities for Arpin’s AT would differ, resulting in a significantly asymmetric binding profile, with site-A3 likely exhibiting stronger binding affinity.

Herein, we report the 1.65-Å crystal structure of the N-terminal globular domain of Arpin, which adopts a unique fold distinct from other endogenous Arp2/3-binding factors. Structural and biophysical analyses confirm that Arpin exhibits the capacity to form homodimers in solution via a conserved interface. Functional assays further establish that Arpin homodimerization is critical for its cooperative inhibition of Arp2/3-dependent actin polymerization, thereby suppressing lamellipodia-driven cell migration and invasion. These structural and functional insights into Arpin’s role as an Arp2/3 complex inhibitor advances our understanding of the molecular mechanism governing cell migration and provides a foundation for designing novel inhibitors targeting cancer metastasis.

## Results

### Overall structure of Arpin globular core

While previous SAXS analysis confirmed a tadpole-like conformation for Arpin^19^, the structural details of its globular core (the "tadpole head") remain unresolved, hindering comprehensive elucidation of its inhibitory mechanism. To address this, we performed crystallographic studies on Arpin orthologs from diverse species (Fig. S1), encompassing full-length constructs and truncated variants (Fig. 1A). Ultimately, a crystal of zebrafish Arpin (residues 9[221) with a single-site mutation C133S (to avoid unnecessary inter-molecular interactions) diffracted to a highest resolution of 1.65 Å. The structure of Arpin was solved by single isomorphous replacement using one Hg derivative and refined against the high-resolution native dataset (see Supporting Information for details; Fig. 1B, Table 1). Within one asymmetric unit, two Arpin molecules form a homodimer exhibiting pseudo two-fold rotational symmetry (Fig. S2A). Notably, residues 187–221 at the C-terminus are missing in the final models of both protomers, which correlates with their intrinsic flexibility and aligns with the SAXS-derived tadpole-like contour. Accordingly, we define the globular core of Arpin as residues 1–186, while the CT of Arpin as residues 187–221.

**Figure 1.**
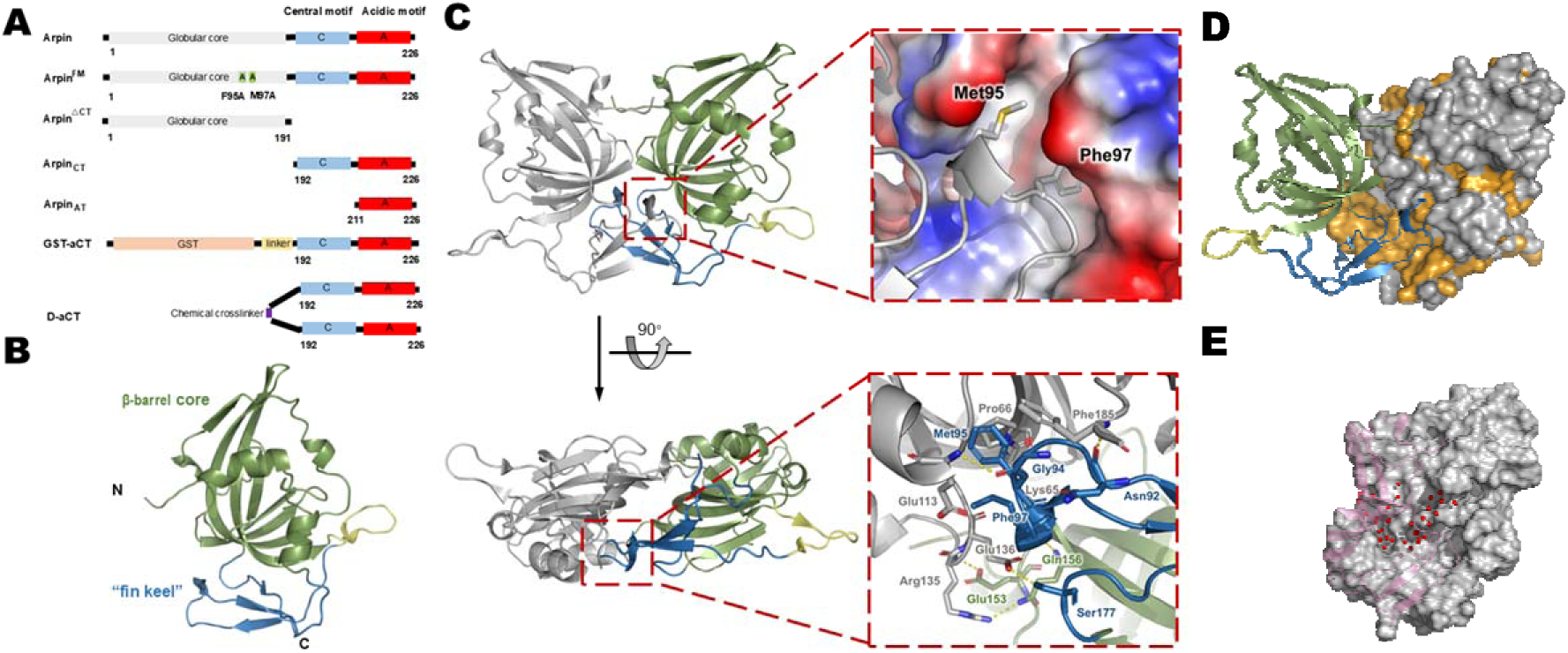
The globular core of Arpin forms homodimers in the crystal. (A). Schematic representation of truncated and mutated Arpin variants. The globular N-terminal domain is shown in grey, the central motif in blue, and the acidic tail in red. (B). Cartoon representation of the globular core of Arpin with β-barrel core (green), “fin keel” (blue) and a short linker (yellow) properly labeled. (C). Close-up view of the dimer interface around contact cluster II. Upper panel: At the tip of the “fin keel”, the hydrophobic side chains of Phe95 and Met97 are shown in sticks. Lower panel: Residues involved in the interactions between adjacent “fin keel” structures are shown in sticks, with inter-protomer hydrogen bonds indicated by dashed lines. (D). Conserved surface patch of Arpin. One protomer of the Arpin homodimer is shown in ribbons, while the other one as surface representation colored by sequence conservation (strictly conserved residues in orange, while others in gray). (E). Water-filled cavity at the Arpin dimer interface. A fully enclosed cavity is formed housing 35 ordered water molecules (in red balls) stabilized by a hydrogen-bond network.

**Table 1.**
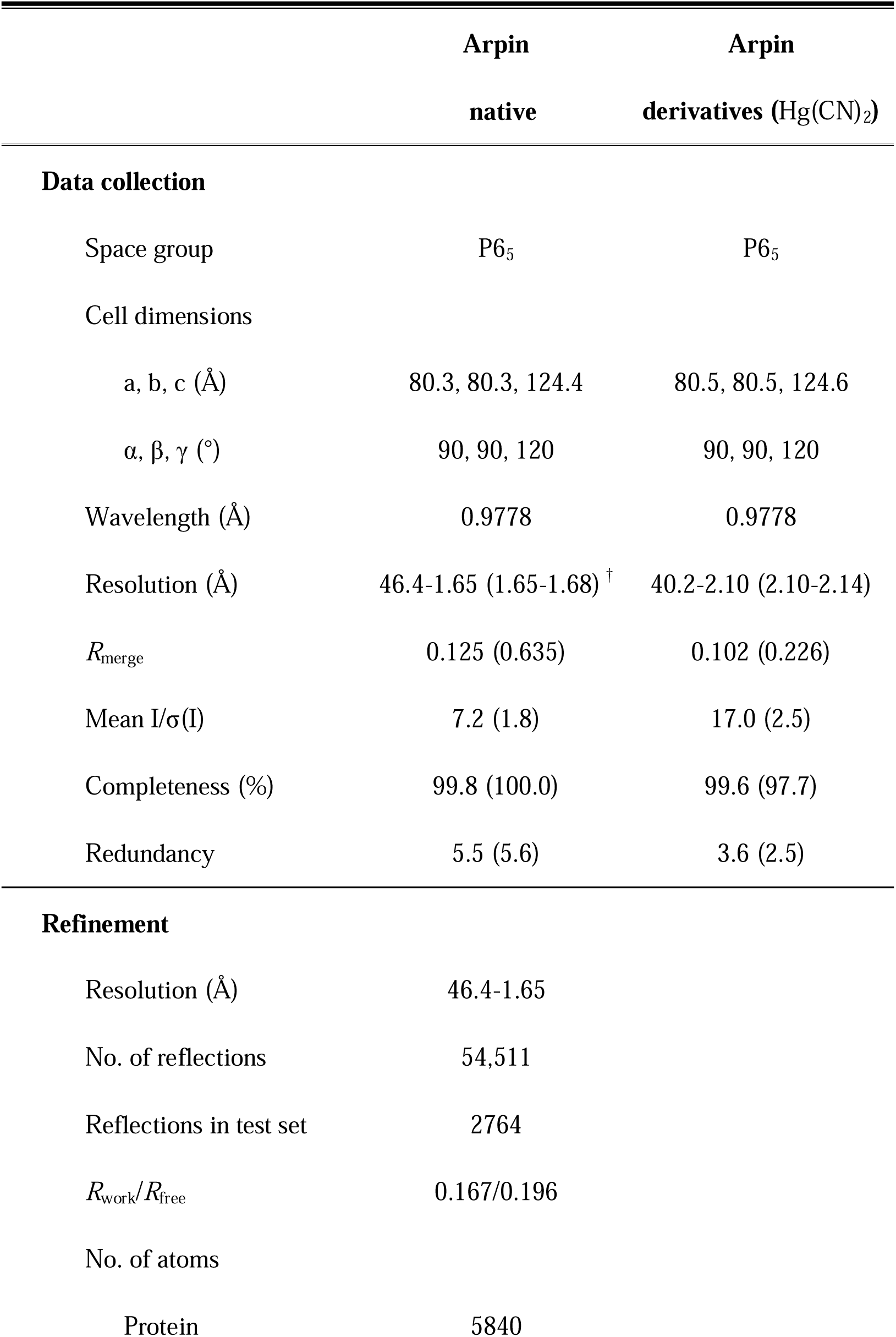
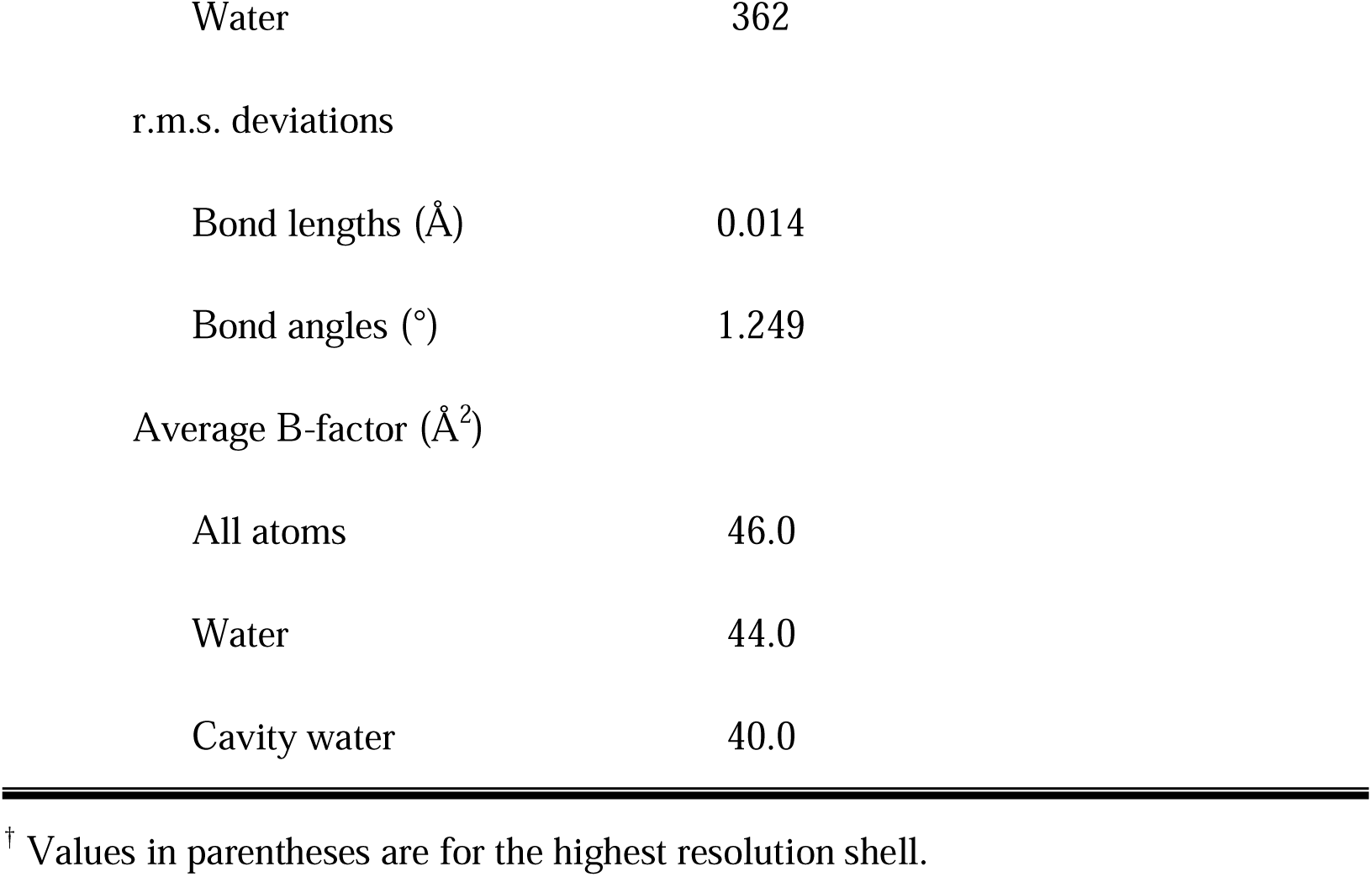
Data collection and refinement statistics.

|  | Arpin<br>native | Arpin<br>derivatives (Hg(CN)<sub>2</sub>) |
| --- | --- | --- |
| **Data collection** |  |  |
| Space group | P6<sub>5</sub> | P6<sub>5</sub> |
| Cell dimensions |  |  |
| a, b, c (Å) | 80.3, 80.3, 124.4 | 80.5, 80.5, 124.6 |
| α, β, γ (°) | 90, 90, 120 | 90, 90, 120 |
| Wavelength (Å) | 0.9778 | 0.9778 |
| Resolution (Å) | 46.4-1.65 (1.65-1.68)<sup>†</sup> | 40.2-2.10 (2.10-2.14) |
| *R*<sub>merge</sub> | 0.125 (0.635) | 0.102 (0.226) |
| Mean I/σ(I) | 7.2 (1.8) | 17.0 (2.5) |
| Completeness (%) | 99.8 (100.0) | 99.6 (97.7) |
| Redundancy | 5.5 (5.6) | 3.6 (2.5) |
| **Refinement** |  |  |
| Resolution (Å) | 46.4-1.65 |  |
| No. of reflections | 54,511 |  |
| Reflections in test set | 2764 |  |
| *R*<sub>work</sub>/*R*<sub>free</sub> | 0.167/0.196 |  |
| No. of atoms |  |  |
| Protein | 5840 |  |

|  |  |
| --- | --- |
| Water | 362 |
| r.m.s. deviations |  |
| Bond lengths (Å) | 0.014 |
| Bond angles (°) | 1.249 |
| Average B-factor (Å<sup>2</sup>) |  |
| All atoms | 46.0 |
| Water | 44.0 |
| Cavity water | 40.0 |

<sup>†</sup> Values in parentheses are for the highest resolution shell.

The overall structure of Arpin’s globular core indeed represents an ellipsoid-shaped domain, characterized by an antiparallel β-barrel framework composed of six β-strands (β1, β2, β3, β8, β9 and β10; Fig. S2B). Among those, the two longest ones, β2 and β3, exhibit pronounced curvature, forming a circumferential band that underpins the barrel’s structural integrity. Encircling this central β-barrel are six helices (α1, α2, η1, η2, η3 and η4) and five additional short β-strands (β4, β5, β6, β7 and β11). The N-terminal 3_10_-helix η1 forms a cap atop the barrel, providing an anchoring point for the N-terminus. Beneath the barrel, three β-strands (β6, β7 and β11) assemble into an extra β-sheet, alongside the 3_10_-helix η3, creating a fin-keel-like structure perpendicular to the barrel’s bottom (Fig. 1B).

Sequence alignments show remarkably low homologies of Arpin with previously known structures (< 20% similarity), yet its architecture adopts the canonical OB-fold, featuring a central β-barrel composed of a highly curved antiparallel β-sheet (Fig. S3A). Structural analysis via DALI server^26^ identifies Arpin as most similar to two OB-fold protein classes (z-score of 6.4): single-stranded DNA-binding proteins (SSBs) from bacteria or yeast mitochondria and metallochaperone proteins in bacterial cation efflux systems. Superimposition studies on Arpin resulted in a 1.95-Å r.m.s. deviation for 71 aligned Cα atoms with *Saccharomyces cerevisiae* Rim1 (PDB entry: 6CQK; Fig. S3B), a mitochondrial SSB, and a 2.41-Å r.m.s. deviation for 68 aligned Cα atoms with *E. coli* CusF (PDB entry: 3E6Z; Fig. S3C), a metallochaperone. Despite the obvious structural parallels, Arpin exhibits distinct functional divergence from SSBs or metallochaperones. It is worth noting that both Rim1 and CusF are significantly smaller than Arpin and lack the N-terminal cap motif and the bottom “fin keel” structure.

### The crystal structure of Arpin unveils homodimeric configuration with extensive intermolecular interactions

In the crystal of Arpin, two protein molecules assemble into a homodimer within one asymmetric unit. The homodimerization of Arpin buries an interface area of 1,722 Å^2^ (16% of the total solvent-accessible surface area), substantially exceeding the other two intermolecular interfaces observed (50 Å^2^ and 187 Å^2^, respectively). The two protomers packed to each other in a two-fold symmetric manner, with their β-barrels approximately parallel to each other, resembling the shape of a butterfly (Fig. 1C). About 20% residues of the Arpin globular core participate in the homodimeric interface, establishing extensive intermolecular interactions, which can be divided into two distinct contact clusters.

Contact cluster I resides between the β-barrels of two Arpin protomers, where the N-terminus of one protomer extends to interact with the 3_10_-helix η4 of the adjacent protomer, establishing predominantly hydrophobic interactions and two inter-protomer hydrogen-bonds (Met8-Asp43 and Glu153-Arg135; Fig. S4). Contact cluster II, formed by the two fin-keel-like structures arranged in a back-to-back manner, stabilizes the projecting structural motifs through five inter-protomer hydrogen-bonds and hydrophobic interactions (Fig. 1C). In particular, at the tip of the “fin keel” (the segment between β6 and β7), the hydrophobic residues Phe95 and Met97 from one protomer protrude their side chains into a hydrophobic pocket constituted by Pro66, Val69, Leu116 and Phe138 of the adjacent protomer (Fig. 1C). Interestingly, the entire “fin keel” region is the most conserved part of Arpin (Fig. 1D & S5), including an absolutely conserved motif ranging residues 90–105 (β6, β7, η3, and connecting loops). The strict conservation of the “fin keel” structure strongly implies that both inter-protomer interactions and the resulting homodimeric configuration are conserved across various species.

Notably, between cluster I and II lies a considerable enclosed cavity (1,628 Å^3^, calculated by MOLE 2.0; Fig. 1E), filled with 35 ordered water molecules (average *B*-factor of 40.0 Å^2^; Table 1) that are partially visible in the experimentally phased electron density (Fig. S6). This cavity is stabilized by an intriguing hydrogen-bond network formed among the ordered water molecules and the hydrophilic side chains at the inter-protomer interface. When the Arpin homodimer is properly formed, its central cavity becomes solvent-inaccessible, preventing water exchange between the cavity and solvent. The tightly restricted mobility of these water molecules further enables their observation in the crystal structure. Unlike classic dimerization interfaces usually dominated by hydrophobic patches, the Arpin homodimer encloses a water-filled cavity, suggesting weak inter-protomer interactions and a dynamic assembly mechanism.

### Dynamic equilibrium between monomeric and dimeric Arpins

The final model of homodimeric Arpin structure encompasses only the globular core, yet the construct for crystallization actually includes the majority of the CT after the globular core. SDS-PAGE analysis of crystallized Arpin confirmed the structural integrity of the CT, with no evidence of proteolytic degradation, indicating that full-length Arpin retains the capacity to adopt a homodimeric configuration. Nevertheless, two unsolved critical questions persist: (1) whether Arpin exists as stable dimers under physiological conditions, and (2) whether full-length human Arpin (hArpin) can also form homodimers as zebrafish Arpin (zArpin) does. To address these questions, we employed size-exclusion chromatography (SEC) and analytical ultracentrifugation (AUC) to characterize the oligomerization behavior of hArpin and zArpin in solution.

Recombinantly expressed hArpin and zArpin both form homodimers in solution (Fig. 2A&B). SEC analysis revealed that hArpin was eluted as a single symmetric peak at 15.6 mL on a Superdex 200 10/300 GL column (bed volume 24mL, GE Healthcare), corresponding to a globular particle with an estimated molecular weight (eMW) of 55.7 kDa—close to the theoretical molecular weight (tMW) of homodimeric hArpin (49.9 kDa). In contrast, zArpin exhibited two SEC elution peaks on the same column: one at 15.8 mL (eMW: 51.8 kDa) and 16.9 mL (eMW: 31.7 kDa), respectively. Given zArpin’s tMW as 25.6 kDa, the two peaks represent the dimeric and monomeric fractions of zArpin, respectively (Fig. 2A). AUC data further clarified their assembly states: hArpin displayed sedimentation coefficients (SC) of 2.330 s (eMW: 18.9 kDa, monomer) and 3.605 s (eMW: 36.4 kDa, dimer), while zArpin predominantly existed as monomers (SC: 2.265 s, eMW: 20.7 kDa) with a minor dimeric fraction (SC: 3.886 s, eMW: 46.5 kDa; Fig. 2B). These results demonstrate that zArpin forms homodimers preferentially at high concentrations (e.g., in concentrated samples prepared for SEC or tightly packed crystalline states), whereas hArpin favors homodimerization in solution.

**Figure 2.**
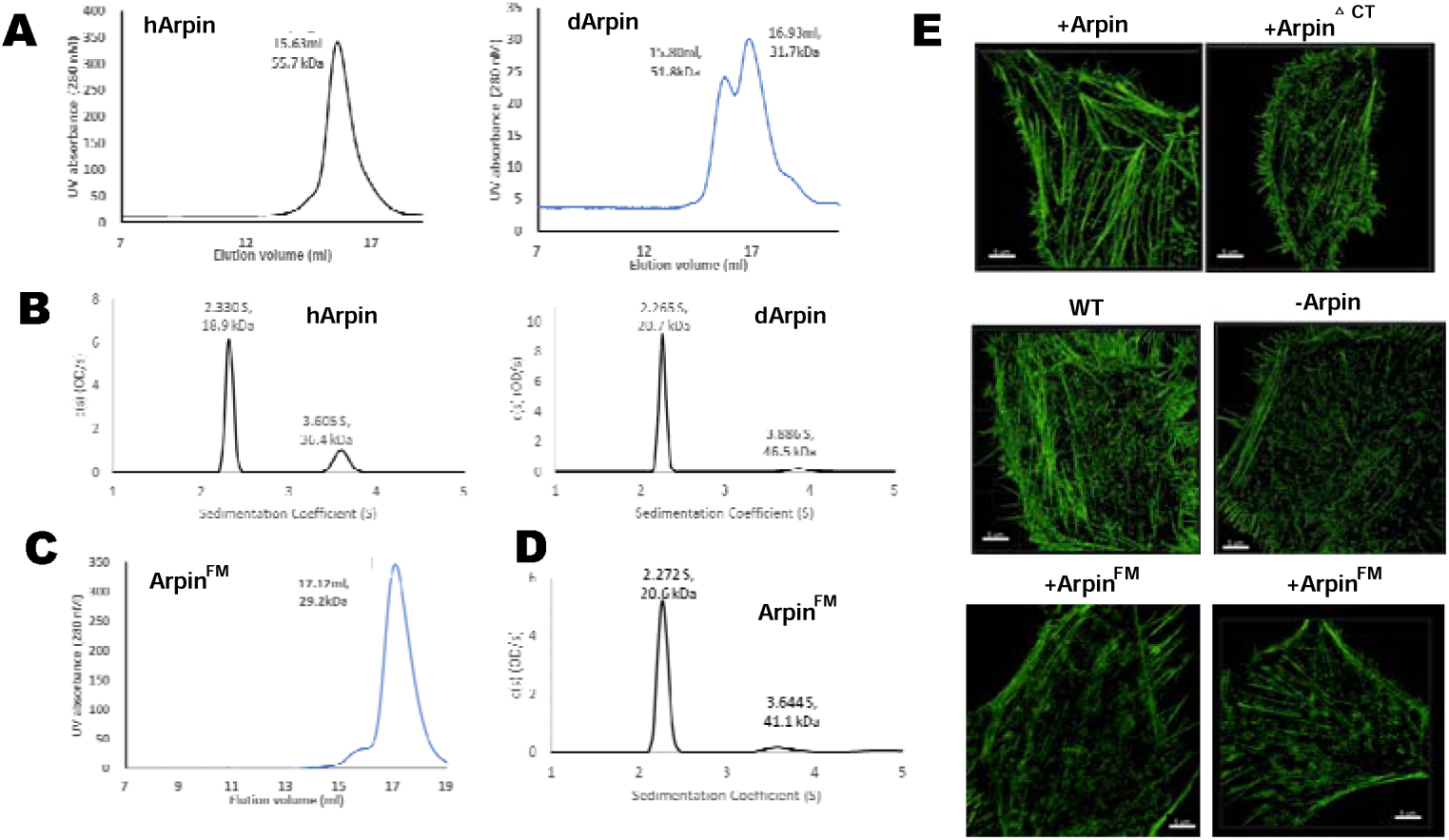
Homodimerization of Arpin regulates the dynamic equilibrium of its oligomeric state and inhibitory function. As demonstrated by (A). SEC and (B). AUC analyses in both human and zebrafish variants, full length Arpin exists in a dynamic equilibrium between monomeric and homodimeric form. (C). SEC and (D). AUC analyses of double-site mutation F95A/M97A. (E). Cytoskeletal staining of MCF-7 cells overexpressing Arpin or its variants. Confocal images of FITC labeled phalloidin-stained MCF7 cells showing the state and distribution of actin filaments (Scale bars, 5 nm). Cells were overexpressed with Arpin, Arpin^ΔCT^ and Arpin^FM^, respectively, as well as knocked down Arpin (-Arpin) and WT cell line.

The homodimeric structure of Arpin unveils the pivotal role of the tightly conserved residues Phe95 and Met97 in stabilizing the inter-protomer interface. Furthermore, Arpin possesses a central enclosed cavity between two adjacent protomers, a structural feature that may foster a higher propensity for dynamic oligomerization compared to typical rigid homodimers. To investigate the functional contribution of Phe95 and Met97 to homodimer stability, we introduced a double-site alanine substitution (F95A/M97A) to hArpin (designated as Arpin^FM^). SEC analysis showed that Arpin^FM^ predominantly adopts a monomeric state in solution (17.12mL, eMW: 29.2kDa; Fig. 2C), a finding corroborated by the result of AUC (SC: 2.272 s, eMW: 20.6 kDa; Fig. 2D). These results strongly suggest that hydrophobic interactions mediated by Phe95 and Met97 are essential for maintaining the structural integrity of Arpin homodimers. Importantly, the substitution does not disrupt the majority of inter-protomer interactions, particularly the hydrogen-bond networks, further underscoring the relative weak stability of Arpin homodimers, as well as its dynamic equilibrium between monomeric and homodimeric states.

### Homodimerization of Arpin enables its cooperative inhibition of Arp2/3 complex

Arpin is known to bind to the Arp2/3 complex and inhibit its nucleation of branched daughter actin filaments, acting as a negative regulator of microfilament network formation^10^. Previous studies have suggested that the Arp2/3 complex contains two distinct binding sites for Arpin, implying the possibility of multivalent interactions^22^. Our structural and biophysical studies demonstrate that Arpin is capable of forming homodimers in solution, raising a speculation that homodimeric Arpin may bind the Arp2/3 complex in a 2:1 stoichiometry, establishing a stable heterotrimer. This manner could enable cooperative inhibition of Arp2/3-mediated nucleation by simultaneously occupying both AT-binding sites. To test this hypothesis, we first assessed the regulation of cytoskeletal organization by Arpin using breast cancer cell line MCF-7 as a model system, as it exhibits remarkably reduced endogenous Arpin expression^12,27^, providing a relevant context for studying Arpin’s regulatory effects.

Cytoskeletal staining revealed that, with overexpressed Arpin, the F-actin filaments in MCF-7 cells appear generally thicker and longer than in control cells (Fig. 2E). On the contrary, MCF-7 cells with Arpin knocked down led to the development of thinner and less extensive actin bundles, particularly in the central region of cells. These observations align with previous studies^10,27^ and demonstrate Arpin’s inhibitory role in suppressing actin filament branching events. In addition, as expected, the overexpression of a truncation lacking the CT (designated as Arpin^ΔCT^, residues 1–191) failed to recapitulate the effects of full-length Arpin, underscoring the fact that Arpin’s binding to Arp2/3 complex is mediated by its CT (Fig. 2E).

The relationship between Arpin’s homodimerization and its inhibitory function had not been previously investigated. To address this, we utilized the double-site mutant Arpin^FM^ that disrupts homodimerization while retaining the overall structure and, in particular, the CT responsible for Arp2/3 binding. Strikingly, MCF-7 cells expressing Arpin^FM^ exhibited a dendritic microfilament structure lacking long bundles, mimicking the effect caused by Arpin^ΔCT^ rather than full-length Arpin (Fig. 2E). Conclusively, Arpin’s ability to inhibit microfilament branching can be impaired by the double-site mutation F95A/M97A. Considering that Arpin^FM^ maintains Arp2/3-binding capability theoretically, these findings strongly suggest the significant role of homodimerization in Arpin’s inhibitory function.

While it is reasonable to infer that Arpin^FM^, which retains the intact CT, can bind to the Arp2/3 complex, this interaction must be experimentally confirmed. Furthermore, if homodimerization plays a key role in the Arp2/3-binding ability of Arpin, the binding affinities of dimeric and monomeric Arpins will be different. To address these issues, we directly evaluate the *in vitro* interactions between the Arp2/3 complex and Arpin variants via surface plasmon resonance (SPR) binding assays, with Arp2/3 immobilized on the sensor chip. During SEC analysis, full-length Arpin samples can be separated into two elution fractions, representing the dimeric and monomeric Arpin, respectively. Both fractions were individually injected into the Arp2/3-coverd chips and exhibited distinct dissociation constant: dimeric Arpin showed a *K*_D_ of 14.1 μM, while monomeric Arpin had a *K*_D_ of 48.9 μM, demonstrate that Arpin homodimerization synergistically enhances its binding affinity to the Arp2/3 complex (Fig. S7, Table 2). In addition, the CT peptide (residues 192–226, designated as Arpin_CT_) was also subjected to SPR assays and displayed a comparable *K*_D_ (17.8 μM) to dimeric Arpin but produced a remarkably higher maximum response value (357.4 RU vs. 94.3 RU), suggesting differences in binding stoichiometry or dynamics. Critically, Arpin^FM^ exhibited a *K*_D_ of 63.1 μM, even weaker compared to monomeric Arpin (Fig. S7, Table 2), indicating that the F95A/M97A double-site mutation disrupts the cooperative binding by preventing homodimerization.

**Table 2.**
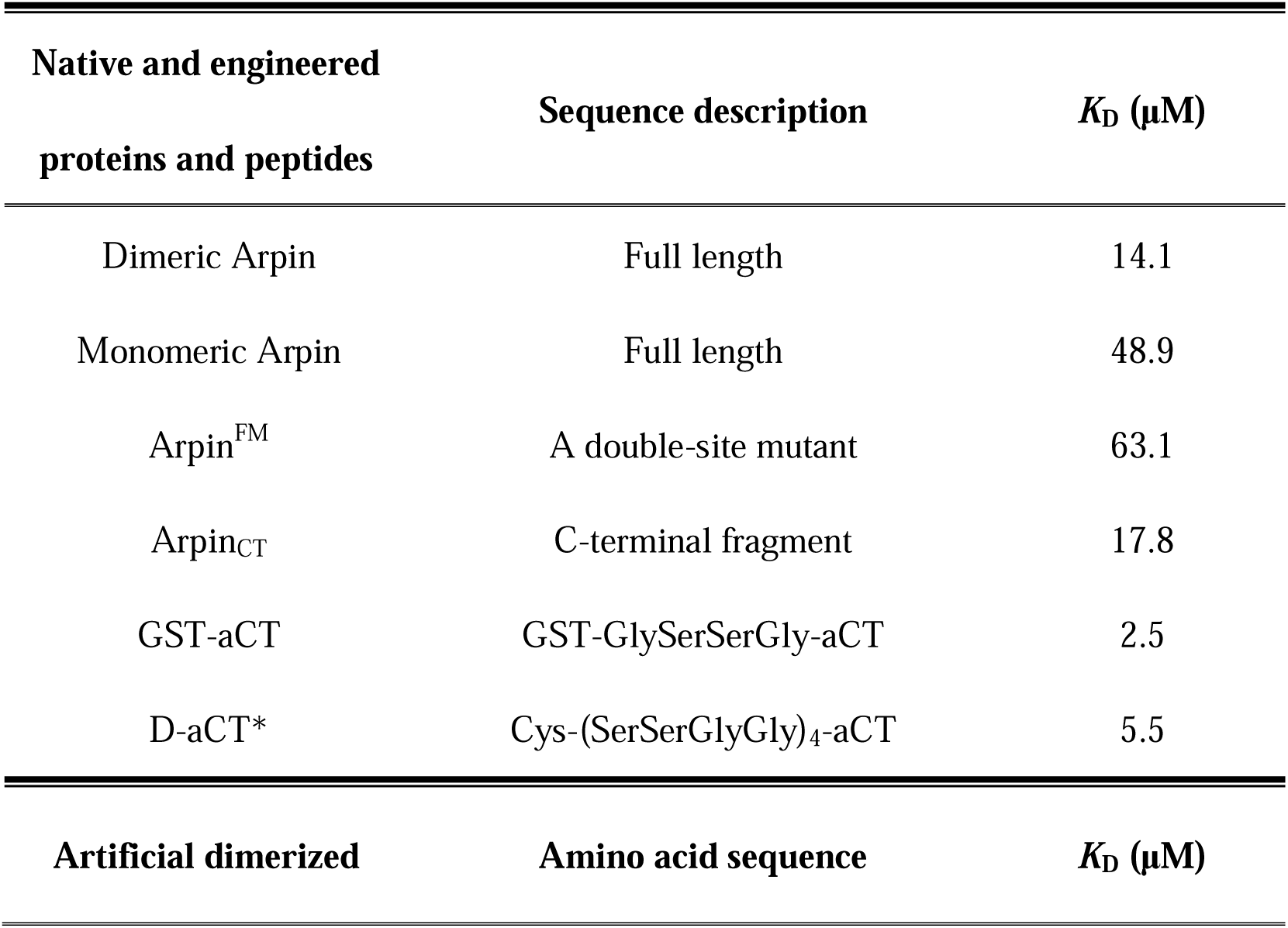
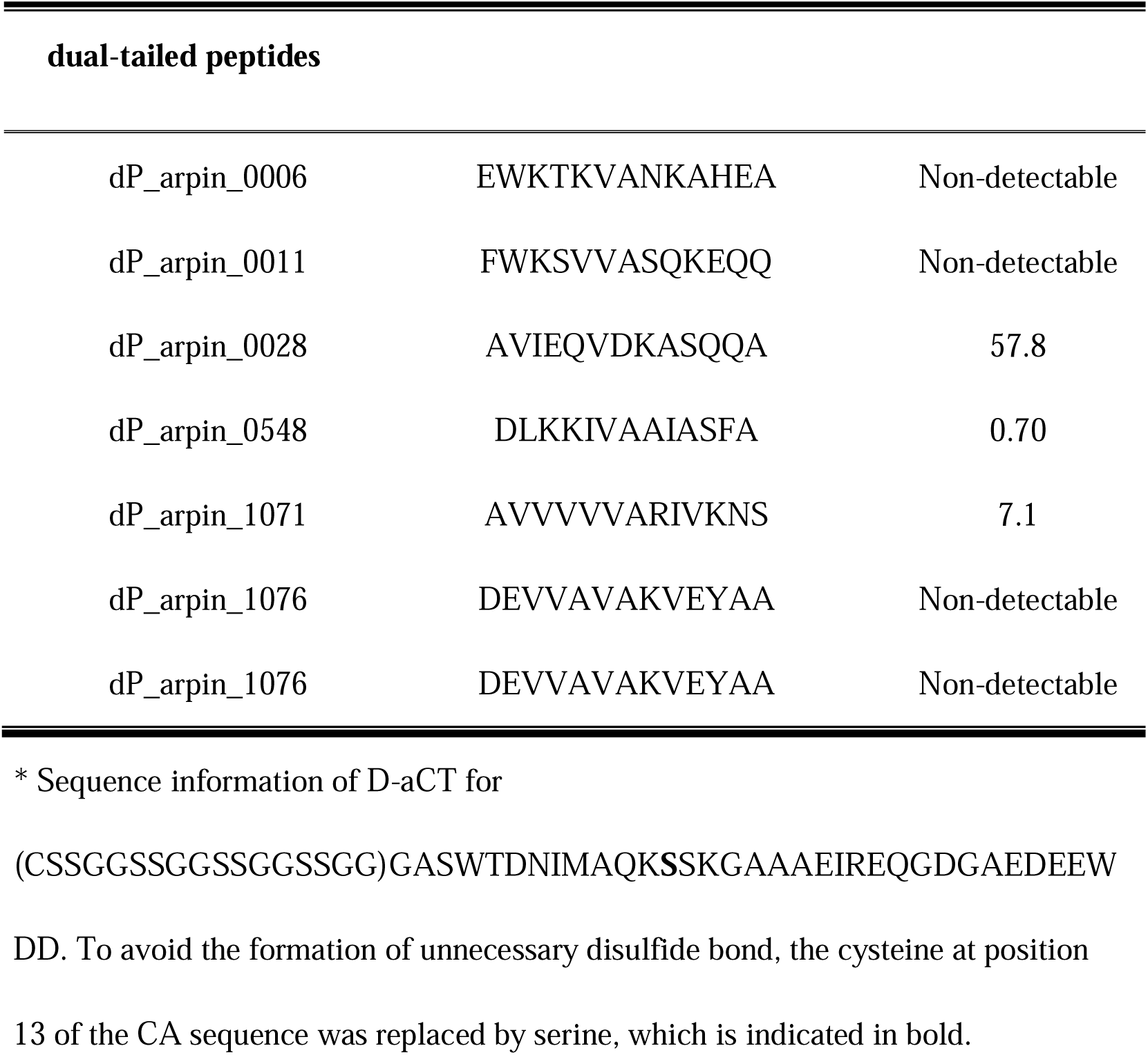
Summary of dimerized peptides and proteins with corresponding SPR binding parameters.

Collectively, these results confirm that Arpin^FM^ retains Arp2/3-binding capacity but adopts a binding pattern resembling the monomeric form. Since Arpin’s CT mediates its association with the Arp2/3 complex while its N-terminal globular domain facilitates its homodimerization, our findings reveal that these structural features are functionally coordinated rather than independent. The intrinsic interplay between Arpin’s homodimerization and Arp2/3 binding establishes a dynamic inhibitory regulatory mechanism, where homodimerization amplifies binding affinity and consequently enhances Arpin’s regulatory function.

### Homodimerization promotes cell steering and opposes directional persistence

Having established the functional interplay between Arpin’s homodimerization and its Arp2/3-binding activity, we proceeded to investigate how this cooperative mechanism influences cell migration. Therefore, we implemented multiple approaches to examine both directional migration capabilities and real-time migratory dynamics in cancer cells expressing Arpin or its engineered variants, including Arpin^FM^, Arpin^ΔCT^, Arpin, and Arpin (residues 211–226, comprising only the minimal inhibitory acidic motif) (Fig. 1A). Still, the breast cancer cell line MCF-7 was selected as the model system.

First, we conducted transwell migration assays. Quantification of migrated cells revealed that Arpin-overexpressing cells exhibited significantly reduced migration (average number: 28.2) compared to control cells (66.6; *P* < 0.0001; Fig. 3A & 3B). In contrast, cells expressing Arpin^ΔCT^ displayed migratory behavior comparable to controls (60.4; *P* > 0.1; Fig. 3A & 3B), indicating that the CT is critical for Arpin’s inhibitory function. The tail-only variants Arpin_AT_ and Arpin_CT_ retained moderate migratory suppression (39.4 and 34.4, respectively, *P* < 0.001 vs. Arpin, Fig. S8), once again confirming that the CT alone is sufficient for partial inhibition of cell migration. Overexpression of Arpin^FM^ indeed led to reduced cell migration, but with significantly weaker inhibitory potency (52.2, *P* < 0.0001 vs. Arpin; Fig. 3A), which is even comparable to Arpin^ΔCT^ (*P* < 0.01; Fig. 3B). Given the fact that Arpin^FM^ contains the CT for Arp2/3 binding but Arpin^ΔCT^ does not, the profound loss of inhibitory activity induced by the F95A/M97A mutation is particularly striking, vividly demonstrating our conclusion that homodimerization is essential for Arpin’s robust function of suppressing cell migrations. In addition, it is noteworthy that, confirmed by western blotting analysis, transfection of full-length Arpin, Arpin^ΔCT^ and Arpin^FM^ led to generally equivalent expression level in the MCF-7 cells for transwell assays (Fig. 3C), ruling out confounding effects of differential protein expression on observed migratory differences.

**Figure 3.**
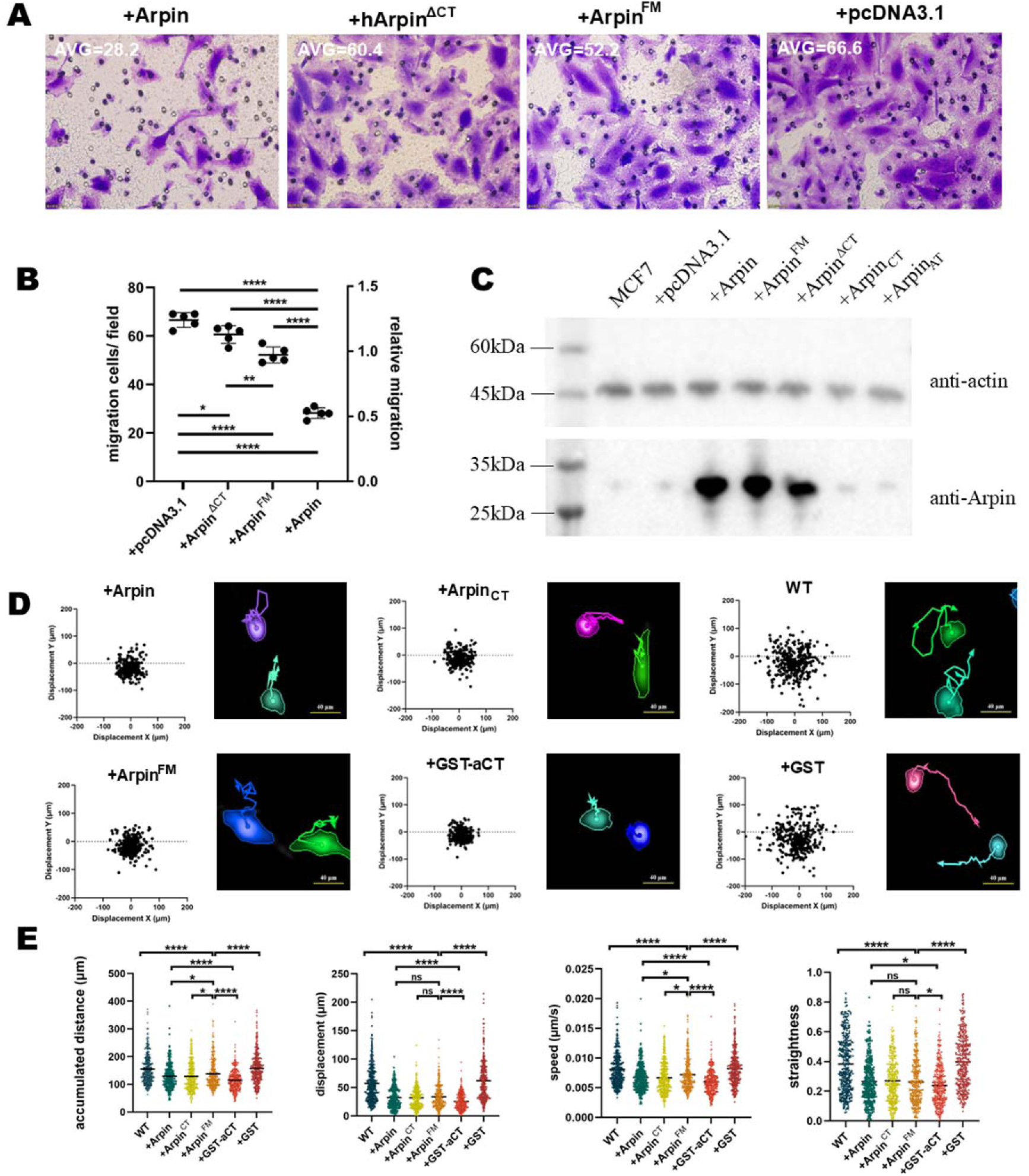
Homodimerization suppresses directional migration by enhancing Arp2/3 inhibition. (A). Transwell assays were performed to analyze the effects of transfection of Arpin with its variants on the migration of MCF-7 cells. Representative images of crystal violet staining of cells captured 24 hours after treatment, original magnification 40×. (B). Numbers of migrated cells through the chamber after 24h. Cells were counted from 10 random microscope fields for each sample in 3 independent experiments. Statistical significance is indicated as follows: *P < 0.05; **P < 0.01; ***P < 0.001; **** p≤0.0001; ns, not significant (P > 0.05). (C). Western blot confirms the expression levels different across Arpin constructs. (D). Representative migration trajectories and vector displacement analysis of the time-lapse tracking of HT1080 cells with stable Arpin expression.HT1080 cells with stable expression of Arpin or control were seeded on collagen-coated dishes and imaged for 6 hours. Each cell’s starting position was set as the origin, and its final position defined the endpoint for vector displacement analysis. Representative migration tracks were also selected and shown to illustrate typical movement patterns. (E) Accumulated distance, net displacement, migration speed, and directionality (straightness) of migrating cells were measured in three independent experiments. Data are presented as mean ± standard deviation. Statistical significance is indicated as follows: *P < 0.05; **P < 0.01; ***P < 0.001; **** p≤0.0001; ns, not significant (P > 0.05).

Subsequently, we investigated the dynamic migratory behavior of HT1080 human fibrosarcoma cells, a highly invasive and metastatic cell line, to further examine the relationship between Arpin’s homodimerization and its inhibitory function. To this end, HT1080 cells expressing Arpin or its engineered variants were seeded at low density in 2D-culture conditions to promote single-cell migration. All the migration trajectories were captured for 6 hours by using time-lapse video microscopy (Fig. 3D, Movie S1) and then quantitatively analyzed across four parameters: accumulated distance, net displacement, speed, and straightness. Compared to controls, the overexpression of full-length Arpin, Arpin_CT_, or Arpin^FM^ significantly reduced net displacement by 44.0%, 44.5%, and 41.4%, respectively (Fig. 3E). As for accumulated distance and speed, cells expressing Arpin, Arpin_CT_, or Arpin^FM^ exhibited reductions of 16.8%, 17.3%, and 11.3%, respectively (Fig. 3E). The apparent discrepancy between the reduction in net displacement and accumulated distance indicates that Arpin barely decreases cell migration speed but promotes cell steering, resulting in reduced directional persistence. These observations corroborate findings from previous studies^10^. Moreover, among the three Arpin constructs evaluated, full-length Arpin and Arpin_CT_ exhibited similar inhibitory capacity (*P* > 0.1; Fig. 3E), whereas Arpin^FM^ showed weaker inhibitory functions than the other two constructs (*P* < 0.1; Fig. 3E). In other words, fusing a monomeric N-terminal domain to the CT even attenuated the CT’s inhibitory function. These results suggest that the evolutionary significance of Arpin’s N-terminal globular domain is inextricably coupled with its homodimerization ability.

### Artificial dimerization confers elevated Arp2/3 complex affinity and impairs cellular mobility

Up to this point in the study, we have shown that the N-terminal globular domain of Arpin underlies its intrinsic homodimerization. Disruption of this dimerization capability leads to markedly reduced Arp2/3-binding affinity and significantly compromises its inhibitory function in regulating directional cellular migration. Building on these findings, we propose the hypothesis that artificial enhancement of Arpin’s dimerization propensity could synergistically augment its suppressive activity. To pursue this objective, we seek a naturally occurring cytosolic protein exhibiting the following attributes: (1) structural and biophysical characterization as a homodimer with enhanced stability relative to Arpin; (2) homodimeric architecture displaying two-fold rotational symmetry; and (3) two C-terminal extensions aligned in the same spatial orientation with inter-terminal distances comparable to those observed in Arpin homodimers.

Ultimately, glutathione S-transferase (GST) was selected to replace the N-terminal globular domain of Arpin, generating a chimeric protein with Arpin’s CT fused to the C-terminus of GST (designated as GST-aCT, Fig. 1A). To ensure the compatibility of GST-aCT with the dual binding sites on the Arp2/3 complex, we performed a structural comparison of Arpin and GST (PDB entry: 6JI6) homodimers. Similar to Arpin, GST forms homodimers with two-fold rotational symmetry, providing a geometric framework that is possible to position the fused CT appropriately on the surface of Arp2/3 complex. Structural analyses showed that the inter-terminal-Cα distances (between the last observable C-terminal residues of two protomers) of Arpin and GST homodimers are 25.0 Å (Fig. S9) and 46.7 Å (Fig. S9B), respectively. To properly accommodate the extended spacing of GST homodimers, we designed three different spacers (-GSSG-, -GSSGSGSTG-, and -KGSGSTSGSG-) for the region between the N-terminal GST and Arpin’s CT, corresponding to theoretical extensions of 15[35 Å. Prior to experimental validation, we initially performed structural modeling of the chimeric GST-aCT and Arp2/3 complex in a 2:1 stoichiometry. Simulations using AlphaFold3^28^ indicated that even the shortest spacer with 4 residues provided sufficient spatial exposure and accessibility of the CT to engage both binding sites on Arp2/3 (Fig. 4A). Based on these results, we selected the spacer -GSSG- for experimental production of engineered GST-aCT (Table 2), as it balanced functional sufficiency with sequence simplicity.

**Figure 4.**
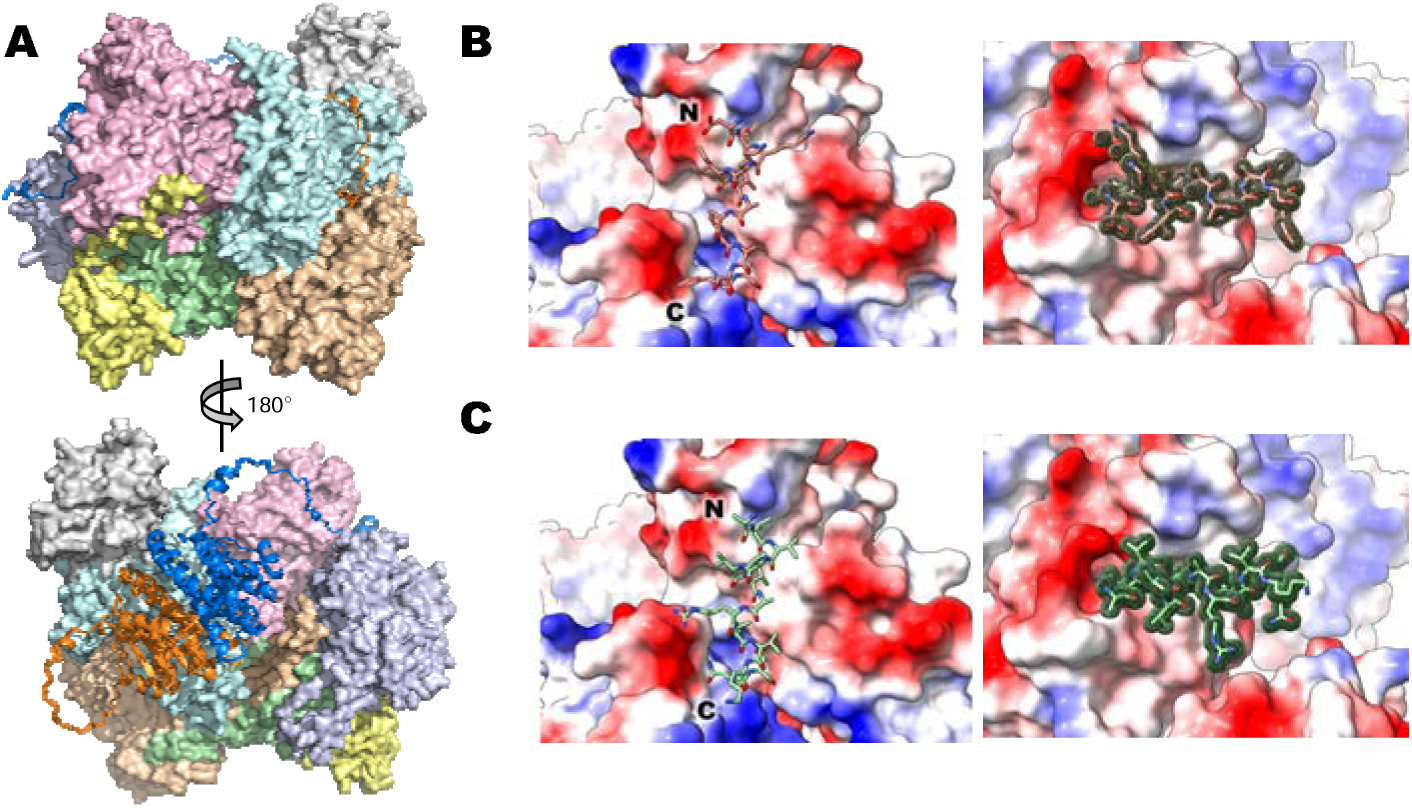
AI-guided engineering of dual-tailed peptides with enhanced affinity for Arp2/3 complex. (A). Structural modeling of GST-aCT bound to the Arp2/3 complex using AlphaFold3. Simulated 2:1 binding stoichiometry confirms that a short linker with 4 residues enable spatial accessibility of both Arp2/3 binding sites. (B) Detailed binding pose of the AI-designed dual-tailed peptide dP_arpin_0548 and its CUTEDGE-generated electron densities. (C) Detailed binding pose of the AI-designed dual-tailed peptide dP_arpin_1071 and its CUTEDGE-generated electron densities..

Recombinantly expressed GST-aCT exhibited a homodimeric state in SEC analysis. The results of SPR assays demonstrated that the engineered GST-aCT possessed a dramatically increased binding affinity to the Arp2/3 complex (*K*_D_ = 2.5 μM; Fig. S7, Table 2), showing 5.6- and 7.1-fold stronger binding compared to Arpin and Arpin_CT_, respectively. These findings indicate that the stable-homodimerization-mediated bivalent binding mode enhances the interaction strength and stability between Arpin’s CT and the Arp2/3 complex.

Encouraged by these results, we investigated whether the reinforced engagement of GST-aCT could further promote the physiological function of CT-binding-mediated inhibition. Live-cell imaging of HT1080 cells expressing GST-aCT exhibited a comprehensive impact on cellular migratory behavior, while overexpression of GST alone exerts no effect, close to the controls (Fig. 3D). Compared to full-length Arpin with dynamic homodimerization, the stably homodimerized GST-aCT led to a 26.3% decrease in accumulated distance and a 55.3% reduction in net displacement, both remarkably stronger than the effects observed with other Arpin constructs in this study (Fig. 3E). Conclusively, our findings demonstrated that while the CT alone exhibited relatively strong inhibitory function, the N-terminal globular domain of Arpin also plays a critical role by mediating homodimerization and further stabilizing the 2:1 Arpin: Arp2/3 complex. And, more importantly, this inhibitory capacity can be further reinforced by the substitution with a more stable N-terminal dimerization domain.

Building on the success of engineered GST-aCT, we next sought to design a minimal inhibitory molecule mimicking homodimerized Arpin. Given that only the CT of Arpin directly interacts with the Arp2/3 complex, we developed a dual-tailed peptide comprising two Arpin’s CTs (D-aCT), which were linked at N-termini by a chemical crosslinker (Fig. 1A). Similar to the design rationale for GST-aCT, a 16-residue spacer (sequence: -(SSGG)_4_-) was incorporated at the N-termini of the CTs to ensure proper engagement with the dual binding sites on the Arp2/3 complex. Due to the presence of arginine and lysine residues in Arpin’s CT segment, which interfere with amine-based crosslinking, we adopted a cysteine-mediated strategy: an exogenous cysteine was added to the N-terminus of CT, while endogenous Cys204 was replaced by serine (Table 2). The peptide was recombinantly expressed and crosslinked using BM(PEG)_3_ (Fig. S10A), which covalently links the N-terminal cysteines (Fig. S10B). Both SEC analysis and mass spectrometry (MS) confirmed that D-aCT formed stable covalent dimers via BM(PEG)_3_, rather than through reducible disulfide bonds. Subsequent SPR assays revealed that D-aCT bind to Arp2/3 with a *K*_D_ of 5.5 μM (Fig. S11, Table 2). This affinity is comparable to that of GST-aCT (2.5 μM) and surpasses that of full-length Arpin or a single Arpin_CT_ peptide (14.1 μM and 17.8 μM, respectively), demonstrating that D-aCT, as a much smaller molecule, achieves robust binding akin to larger Arpin-based constructs.

### AI-aided design of the dual-tailed peptide leads to further enhanced binding affinity for Arp2/3

To further enhance D-aCT’s binding affinity to the Arp2/3 complex, we conducted targeted peptide design via CUTEDGE (Chain-free Underinformed Targeted Electron Density Generative Explorer), a 3D diffusion model operating in electron density space. This deep generative framework, which we have developed for protein/peptide binder design (unpublished), enables rapid generation of structurally plausible binders with high affinity to a given target protein.

Previous studies have identified two distinct binding sites for Arpin on the Arp2/3 complex: site-A3 and site-A2C1^22,23^. Arpin’s CT engages these sites through divergent docking modes, thereby introducing structural complexity that poses significant challenges for computational design approaches. Conventional methods, which typically focus on single-target interactions during the generative process, are ill-suited to handle the dual-interface requirement, necessitating advanced modeling capabilities beyond standard design paradigms.

Given current limitations in simultaneous multi-target binder design, we employed a staged computational design strategy by designing the peptide against one binding site and evaluating the designs at both sites. Among the two Arpin-binding sites, only site-A3 has been confirmed by experimental structures^23^, whereas site-A2C1 is a putative one based on the 2:1 NPF: Arp2/3 complex structures^20,21^. We selected the cryo-EM structure of Arpin_CT_: Arp2/3 (PDB entry: 7JPN, 3.2 Å) as the target model for designing, which displays a bound Arpin_CT_ at its site-A3, with its central domain in helical conformation and its AT in disordered conformation^23^. Considering the conservation of acidic motif across various modulators of Arp2/3, we speculated that further optimization of the AT segment to achieve stronger binding affinity would be highly challenging. Therefore, we focused on designing a 13-residue α-helix, matching the helical central domain of Arpin_CT_ (residues 194 - 206). Within 4.5 hours, CUTEDGE generated 600 electron density maps representing helical peptides bound to the target region. Subsequent iterative refinement yielded 2,292 sets of structural coordinates that fit the generated electron densities.

By anchoring our designs to the experimentally resolved site-A3, we ensured the functional relevance of designed helical peptides while exploring novel sequence space. However, computational evaluation of both binding sites was necessary before experimental assessment. To address the lack of structural details for the site-A2C1, we constructed a homology-modeled of human Arpin_CT_: Arp2/3 complex in a 2:1 stoichiometry based on the cryo-EM structure of the 2:1 NPF: Arp2/3 complex. As a result, one of the two Arpin_CT_ was appropriately positioned for engagement with site-A2C1 (Fig. S12). Rigid-body docking simulations were then performed to assess the binding modes of designed peptides. A multi-parameter scoring pipeline integrated geometric metrics (hydrogen bond networks, salt bridge interactions, steric clashes, and buried solvent-accessible surface areas) and energetic calculations (absolute binding free energies via Adaptive Poisson-Boltzmann Solver, APBS) to prioritize candidates (partial sequences with high scores are listed in Table S1 & S2). This framework identified six structurally viable peptides with predicted binding free energies comparable to native Arpin interactions.

To experimentally validate the AI-designed peptides, the central domain of the CT regions in D-aCT were substituted with the six virtually screened sequences, labeled according to their design IDs (Table 2). The chimeric peptides were recombinantly expressed and crosslinked using the same protocol as D-aCT production. Subsequent SPR assays with the human Arp2/3 complex revealed further enhanced binding affinities for two designed dual-tailed peptides: dP_arpin_0548 (*K*_D_ = 0.70 μM; Fig. S11) and dP_arpin_1071 (*K*_D_ = 5.51 μM; Fig. S11). In fact, dual-tailed dP_arpin_0548 exhibited a 20-fold improvement in binding affinity compared to homodimeric Arpin and a 7.9-fold increase over D-aCT, demonstrating that scaffold-constrained design via deep generative models can achieve dual-interface compatibility and a further enhanced binding affinity of the 2:1 Arpin: Arp2/3 complex.

## Discussion

The metastatic and invasive hallmarks of cancer have been regarded as formidable barriers to the overcoming of cancer recurrence and therapeutic resistance. Although accumulating evidence identifies Arpin as a key regulator of cancer metastasis through its modulation of cell migration, the molecular mechanisms underlying its function remain poorly understood, hindering the development of targeted therapies aimed at inhibiting tumor dissemination. Our work presents the first high-resolution structural characterization of Arpin, addressing a critical gap in the mechanistic understanding of this protein. Importantly, we identified, for the first time, that Arpin can adopt a homodimeric conformation mediated by its N-terminal domain in solution. This discovery provides a novel structural framework to resolve the aforementioned contradictions: Arpin’s interaction with the Arp2/3 complex is not a static 2:1 binding mode but rather a dynamic, positively cooperative binding process. Functional assays further revealed that disrupting Arpin’s homodimerization capacity via F95A/M97A double substitutions resulted in a mutant protein with significantly reduced migratory inhibitory activity compared to full-length Arpin or the isolated CT fragment alone. These findings underscore that dimerization is indispensable for Arpin’s suppressive activity, suggesting that its cooperative assembly may serve as a critical regulatory node in controlling metastatic progression.

Positive cooperativity is a prevalent phenomenon in homo-oligomeric proteins, exemplified by the oxygen-binding mechanisms of hemoglobin and myoglobin. In hemoglobin, a tetrameric protein, oxygen binding to one subunit induces conformational changes that enhance the affinity of subsequent subunits for oxygen, a hallmark of positive cooperativity. This cooperative mechanism relies on a strict 1:1 stoichiometric relationship between subunits and ligands, with each subunit independently contributing a consistent binding surface. In contrast, the interaction between Arpin and the Arp2/3 complex deviates from a rigid 1:1 stoichiometry, instead exhibiting a dynamic one. This behavior arises from Arpin’s N-terminal globular domain-driven homodimerization, which enables cooperative engagement of two structurally and functionally asymmetric binding sites: site-A3 and site-A2C1. Site-A3 likely exhibits higher affinity, dominating under low-concentration conditions (e.g., ITC assays and cryo-EM samples) to yield a 1:1 stoichiometry. In contrast, at elevated concentrations (e.g., concentrated samples for SEC analysis), Arpin occupies both sites, adopting a 2:1 configuration. Physiologically, localized concentration effects via a regulated recruitment mechanism of Arpin likely facilitate the 2:1 mode. Similar recruitment has been found in the formation of other complexes, like PD-L1 and BMS forming 1:1 complex first, and then recruits a second PD-L1 to form a dimer, providing dual binding sites for the initially bound BMS^29–31^.

Moreover, Arpin’s dynamic modulation could be further influenced by the inherent structural complexity and flexibility of the Arp2/3 complex, coupled with its interplay with actin filaments and other activation factors in the cellular milieu. To date, the Arp2/3 complexes subjected in structural characterizations with Arpins were isolated inactive form, where the Arp2 and Arp3 subunits remain spatially separated. Upon activation, the complex undergoes significant conformational rearrangements that position Arp2 and Arp3 at the branch tip to nucleate nascent actin filaments^32–34^. Additionally, NPFs recruit actin subunits to the barbed end of the Arps to form the initial polymerization seed and promote all seven subunits of Arp2/3 complex binding to the side of pre-existing mother filament^32,35–38^. Yet up to now, no structural studies of the Arpin: Arp2/3 complex have taken actin into consideration.

Based on our findings, we propose the “butterfly” model to elucidate the assembly logic of Arpin-Arp2/3 interactions. The crystal structure of the Arpin homodimer reveals two protomers resembling the “wings” of a butterfly, with their missing CT segments extending outward as flexible “cerci” (Fig. 5A). The model centers on a two-step binding process. First, initial preferential occupation of site-A3 by the CT of a single Arpin molecule leads to the formation of a stable 1:1 complex, which inhibits the short-pitch transition of NPF binding by locking the inhibitory tail of Arp3 at the high-affinity site-A2C1^8,20^, and competes with NPF for the low-affinity binding site on Arp3^23^. Second, a subsequent binding pathway of the second Arpin molecule could display dynamic preference (Fig. 5B). One possibility involves directly engaging Arpin’s CT with site-A2C1, albeit with weaker affinity. Spatial effects then promote proximity between the two Arpin molecules, facilitating homodimerization[[two “wings” assembling into a complete “butterfly”. This dimerization stabilizes the complex via interactions at the unstable homodimer interface and low-affinity site-A2C1, collectively forming a stable oligomer with three distinct interaction sites. However, experimental evidence strongly supports a more efficient pathway: the second Arpin molecule first associates with the pre-bound Arpin to form a homodimer. This intact “butterfly” spatially reorients the second “cercus” toward site-A2C1, significantly boosting CT-binding efficiency at site-A2C1 and stabilizing a structurally robust oligomer. Notably, as a more stable homodimer, engineered GST-aCT recapitulates this pathway, exhibiting superior Arp2/3 binding compared to monomeric Arpin and enhanced inhibition of cellular motility.

**Figure 5:**
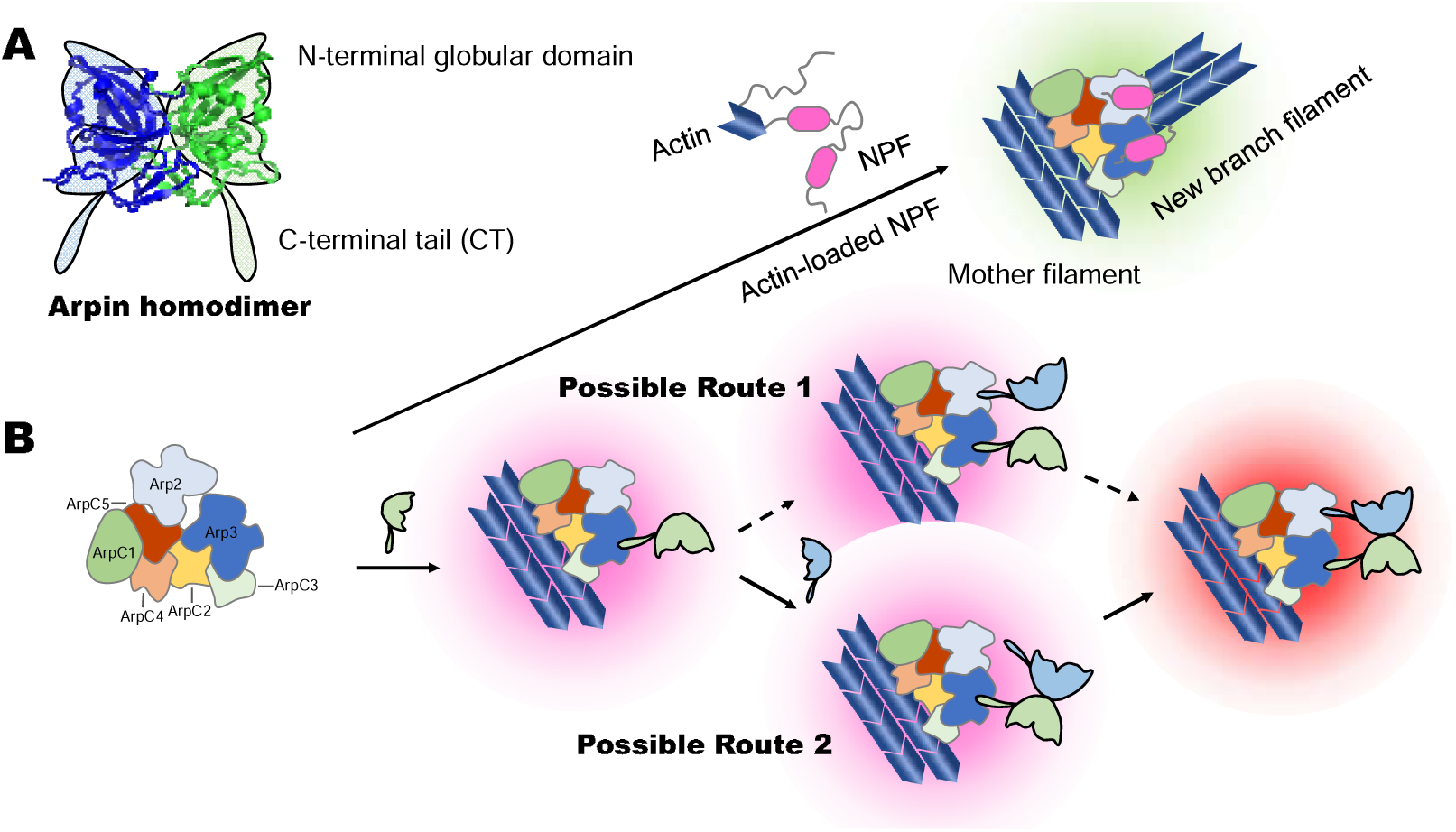
Proposed “butterfly” model illustrates Arpin–Arp2/3 assembly and inhibitory mechanisms. (A). The shape of Arpin’s homodimeric conformation resembling a butterfly. The two globular domains form the “wings”, while the unobserved flexible CT are conceptualized as “cerci.” (B). Schematic model of two potential inhibitory pathways. Upper panel: In the “green light” mode, Arp2/3 binds NPF and actin to promote microfilament branching. Lower panel: In the “red light” mode, Arpin competitively occupies the Arp2/3 binding sites through two possible routes: (1) sequential binding at site-A3 and site-A2C1 followed by dimerization, or (2) initial homodimerization followed by cooperative dual-site engagement.

Regarding the positive cooperativity proposed in the butterfly model, further experimental validation is required. One potential approach involves employing *in situ* EM methods for structural characterization of Arpin: Arp2/3 complexes bound to actin filaments. Nonetheless, the low molecular weight of Arpin poses significant technical challenges for such studies. Alternatively, stabilized GST-aCT homodimers could be directly utilized for structural analysis of the complex. This methodology is precedented by studies of the GST-Crn1:Arp2/3 complex, which successfully resolved its 2:1 binding configuration^22^. Another complementary strategy entails preparing stabilized Arpin homodimers via chemical crosslinking, followed by structural characterization of the resultant 2:1 Arpin: Arp2/3 complexes. Additionally, experimental investigation of the asymmetry in positive cooperativity requires site-directed mutagenesis of the two binding sites on Arp2/3. This could be achieved through knockout/knock-in strategies to generate Arp2/3 complexes with mutations at individual binding sites, enabling verification of the dynamic binding order between the two sites and the determination of independent binding affinity measurements unaffected by cooperative effects. Collectively, these approaches aim to dissect the molecular basis of cooperative binding and validate the mechanisms proposed in the butterfly model.

The potent inhibitory activity of the engineered dimeric GST-aCT construct inspired us to develop molecules with enhanced inhibitory potency, aiming to explore the feasibility of targeting cancer metastasis with therapeutic agents. To this end, we first simplified the molecular form by designing a dual-tailed peptide, D-aCT, which retains two CT segments and shows Arp2/3-binding affinity comparable to that of GST-aCT. Subsequently, using our previously developed deep generative model for protein design (CUTEDGE), we re-engineered the central domain of the CT module and successfully generated dP_arpin_0548, which exhibits the highest Arp2/3-binding affinity observed in this study. Next, both D-aCT and dP_arpin_0548 require validation of their actual effects on cancer metastasis, necessitating intracellular delivery of the peptides. Potential strategies involve conjugating cell-penetrating peptides to facilitate peptide uptake or utilizing liposome nanoparticle delivery systems to encapsulate and deliver the engineered peptides into cells. These approaches aim to bridge the gap between molecular function and therapeutic application while addressing the practical challenges of intracellular targeting and efficacy.

This study reveals a critical interplay between Arpin homodimerization and its inhibitory function in regulating cell migration, and demonstrates a pronounced positive cooperative effect mediated by homodimerization. However, two key questions remain to be addressed. First, why does Arpin form dynamic homodimers instead of stable ones, and what is the significance of this dynamic behavior in regulating its inhibitory activity? A potential regulatory mechanism may involve as-yet-unidentified factors that modulate Arpin’s activity through disruption of its homodimerization, rather than through direct competition for the AT-binding site. Second, what underlies Arpin’s intriguing inhibitory mechanisms, which minimally reduce migration velocity yet paradoxically enhance steering capacity? Steering depends on *de novo* lamellipodia formation via actin branching networks, whereas directional persistence relies on the stabilization of existing networks. Our data indicate that Arpin’s function does not directly inhibit the formation of new branches but instead facilitates the disassembly of pre-existing networks or prevents rebranching during their turnover. Further investigation into the molecular mechanism of Arpin’s homodimerization-dependent inhibition, coupled with its dynamic regulation, may pave the way for developing Arpin-based therapeutic strategies to combat cancer metastasis.

## Supporting information

Supporting materials and methods, Tables S1 to S2, Figures S1 to S12 and Movies Legends for S1

## Methods

### Protein expression and purification

The human and zebrafish Arpin genes (*Homo sapiens*, NM_182616, *Danio rerio*, NM_001017780, including truncations and mutants) were synthesized (Sangon Biotech, China) and inserted into expression vector pET-32a. The fusion protein was overexpressed in *Escherichia coli* BL21. The F95A and M97A mutation was introduced by double mutant site-direct mutagenesis and verified by sequencing. Cells were grown to OD_600_≈0.8 in LB media supplemented with 1 µg/ml ampicillin at 37°C and were induced by 1 mM IPTG for 16 hours at 4°C for expression. Cells were harvested by centrifugation and lysed by high pressure homogenizer for twice after resuspended in buffer containing 50mM Tris-HCl pH 8.0, 200mM NaCl. The fusion protein was isolated from the soluble cell lysate by Ni-affinity chromatography. After using PreScission Protease to cleaves the expression tag, Arpin was purified by Hitrap Q HP column (GE Healthcare Biosciences, USA) for anion-exchange chromatography and a Superdex-200 gel filtration column (GE Healthcare Biosciences, USA). The purified protein was concentrated to 25mg/ml for crystallization or storage at -80°C.

### Crystallization

Zebrafish Arpin (a.a. 10-221) with C133S mutation was crystallized at 16°C using hanging drop-diffusion methods. 1μl of protein solution (10 mg/ml in buffer of 50mM Tris-HCl pH 8.0, 200mM NaCl) and 1μl of reservoir solution (1.6 M Sodium citrate pH6.5) were mixed. The droplets were set on a siliconized glass sheet and inverted on the chamber with 1ml of reservoir solution and then sealed with vacuum silicone grease. Crystals formed after four days. The crystals were flash-frozen by liquid nitrogen for further data collection. Heavy atom derivatives were obtained by soaking the crystals of Arpin in the mother liquor containing 10 mM of Hg (CN)_2_ for 7 min for phasing analysis.

### Data Collection, Structure Determination, and Refinement

The native data sets and heavy atom derivative data sets were all collected at beamline BL-18U1 of Shanghai Synchrotron Radiation Facility with wavelength of 0.9778 Å for the Arpin crystals. The crystals were kept at 100 K during X-ray diffraction data collection. Data were indexed and scaled with HKL2000. Phases were solved by single-wavelength anomalous dispersion method using PHENIX. autosol. All refinement procedures were carried out with PHENIX. refine and COOT. Table 1 shows the detailed statistics of data collection and refinement.

### Surface plasmon resonance

All SPR experiments were performed using Biacore 8K instruments (GE Healthcare) at 25°C, with PBS containing 0.05% (v/v) Tween-20 as the running buffer. Ligands were immobilized on CM5 sensor chips via standard amine coupling, following the manufacturer’s instructions (GE Healthcare). Kinetic parameters were derived by fitting the sensorgrams to fit binding model. Increasing concentrations of Arpin variants were injected at a flow rate of 30μL/min, with a 60s association phase followed by a 60s dissociation phase. For Arp2/3 complex binding assays, dimeric and monomeric fractoins of Arpin, as well as Arpin^FM^, were tested in a concentration range of 3.125–200μM, while Arpin_CT_, GST-aCT and D-aCT were tested at 0.625–40μM. AI-designed Arpin variants were tested in a concentration range of 0.156–10μM. Capture levels were optimized to achieve appropriate analyte responses. The anti-tag capture antibody was covalently coupled to the CM5 chip via amine coupling and regenerated according to the manufacturer’s protocol.

### Cell culture and transfection

The human breast cancer cell line MCF7 and fibrosarcoma cell line HT1080 were cultured in high-glucose DMEM without glutamine, supplemented with 10% FBS, at 37°C, 5% CO_2_. Mammalian codon-optimized synthetic human Arpin genes (including full-length, mutant, and truncated variants) were cloned into the pcDNA3.1 expression vector (Tsingke, China). Duplex siRNAs were synthesized as follows: Arpin knock down-specific sequence, 5 ′ -GGCGCUAGUUGGACCGAUA-3′(sense) (Sangon Biotech, China). Transient and lentiviral transfections were performed using Lipofectamine 2000 (Invitrogen, USA) according to the manufacturer’s protocol.

### Cytoskeletal staining assay

MCF7 cells grown on glass coverslips were fixed with 4% paraformaldehyde in PBS for 10 minutes at room temperature, followed by three washes with PBS. The cells were then permeabilized with 0.5% Triton X-100 in PBS for 5 minutes. F-actin was stained using TRITC-conjugated phalloidin (Yeasen, China) by incubating the samples in the dark for 30 min. After additional PBS washes, the coverslips were mounted in glycerol mounting medium. Fluorescence imaging of MCF7 cells was performed using structured illumination microscopy (SIM) on a DeltaVision OMX V3 system (GE Healthcare). SIM image stacks were acquired with serial Z-sections at 125nm intervals and reconstructed using softWoRx 6.1.1 (GE Healthcare).

### Transwell assay

For the transwell migration assay, 300[μL of FBS-free medium containing 2 ×10^4^ MCF7 cells was added to the upper chamber of a transwell insert (#3422, Corning, USA), while 750μL of DMEM supplemented with 10% FBS was placed in the lower chamber. After incubation for 24h at 37°C under 5% CO_2_, cells were fixed with 4% paraformaldehyde for 20 min. Migrated cells on the lower surface of the membrane were stained with 0.1% crystal violet for 20 min at room temperature. Non-migrated cells on the upper surface were gently removed using a cotton swab. Images were acquired from three randomly selected fields using an Olympus FV3000RS confocal microscope.

### Live-cell migration assay

HT1080 cells stably expressing mScarlet-tagged Arpin variants were seeded at low density on glass-bottom dishes coated with fibronectin and allowed to attach in complete DMEM. Cell migration was monitored using the PE Operetta High Content Screening System (PerkinElmer, USA) at 37°C and 5% CO_2_ for 6h. The instrument automatically scanned (×10) with 9 regions of cell images throughout each well on the plate and images were captured every 10 min. Time-lapse data were analyzed using Harmony 4.0 software (PerkinElmer, USA).

### Crosslinking

To construct the crosslinked D-aCT peptide, Arpin’s CT with an N-terminal Cys and a following spacer (-[SSGG]_4_-) was recombinantly expressed in *E. coli* (Table 2). The expressed peptide possesses additional N-terminal His-tag and SUMO-tag, enabling the purification via Ni-affinity chromatography. Subsequently, the additional tags were removed by overnight Ulp1 cleavage. The cleaved peptides were further purified by ion exchange chromatography without reducing agent. Subsequently, the purified peptides were incubated at room temperature with BM(PEG)_3_ (1,11-bis(maleimido)triethylene glycol) at a molar ratio of 1:1.3 for 30 minutes. After incubation, the crosslinked fraction was isolated from the intact fraction by SEC via a Superdex 75 column (GE Healthcare).

### Western Blot

MCF7 cell lysates were prepared for protein samples, separated by SDS-PAGE (40μg per lane), followed by transfer onto PVDF membranes. After blocking with 5% non-fat milk in TBST, the membranes were incubated with primary antibodies against Arpin (Cusabio, China) and actin (Abmart, China). Subsequently, membranes were incubated with appropriate HRP-conjugated secondary antibodies: goat anti-rabbit (Cell Signaling Technology, USA) or goat anti-mouse (Abmart, China). Protein signals were detected using the Tanon 5200 Multi-imaging System (Tanon, China) and quantified with ImageJ software.

### Accession numbers

Coordinates and structure factors have been deposited in the Protein Data Bank with accession number 6JCP for crystal structure of zebrafish Arpin.

## Supplementary information

This manuscript includes 3 supplementary files that provide detailed experimental procedures and additional experimental results not included in the main text: Supplementary Information (SI): Contains comprehensive descriptions of experimental details and supplementary experiments that support the findings presented in the main manuscript.

Extended Data: Tables S1 to S2, Figures S1 to S12 and Movies Legends for S1. Extended Data_Movie_S1.

## Acknowledgements

We are very grateful to the staff of the Structural Biology Core facility (Institute of Biophysics, Chinese Academy of Sciences) for their technical assistance, especially to Ms. Ya Wang, Ms. Xiaoxia Yu, Ms. Yuanyuan Chen and Mr. Yi Han.

We would like to thank Ms. Shuoguo Li and Miss. Yun Feng from the Center for Biological Imaging (CBI, Institute of Biophysics, Chinese Academy of Science) for their help of taking and analyzing SIM images. We would also like to thank the Biological Reaction Facility (Ms. Wenjuan Zhang) and the Imaging Facility (Mr. Jun Chen and Miss. Chunhua Zhang) of the National Center for Protein Sciences Beijing for their assistance. This work was supported by the National Natural Science Foundation of China (Project No. 31871287, 31400638 and 31470792), and Beijing Municipal Science & Technology Commission grant [Z221100003522020].

## Notes

### Competing Interest Statement

The authors have declared no competing interest.

### Summary of Updates

This manuscript has updated the author informations.

## References

1 Yamaguchi, H. & Condeelis, J. Regulation of the actin cytoskeleton in cancer cell migration and invasion. Biochimica et Biophysica Acta (BBA)-Molecular Cell Research 1773, 642–652 (2007).

2 Ridley, A. J. et al. Cell migration: integrating signals from front to back. Science 302, 1704–1709 (2003).

3 Lai, F. P. et al. Arp2/3 complex interactions and actin network turnover in lamellipodia. The EMBO journal 27, 982–992 (2008).

4 Konietzny, A., Bär, J. & Mikhaylova, M. Dendritic actin cytoskeleton: structure, functions, and regulations. Frontiers in cellular neuroscience 11, 147 (2017).

5 Goley, E. D. et al. An actin-filament-binding interface on the Arp2/3 complex is critical for nucleation and branch stability. Proceedings of the National Academy of Sciences 107, 8159–8164 (2010).

6 Goley, E. D. & Welch, M. D. The ARP2/3 complex: an actin nucleator comes of age. Nature reviews Molecular cell biology 7, 713–726 (2006).

7 Suraneni, P. et al. The Arp2/3 complex is required for lamellipodia extension and directional fibroblast cell migration. Journal of Cell Biology 197, 239–251 (2012).

8 Shaaban, M., Chowdhury, S. & Nolen, B. J. Cryo-EM reveals the transition of Arp2/3 complex from inactive to nucleation-competent state. Nature structural & molecular biology 27, 1009–1016 (2020).

9 Alekhina, O., Burstein, E. & Billadeau, D. D. Cellular functions of WASP family proteins at a glance. Journal of cell science 130, 2235–2241 (2017).

10 Dang, I. et al. Inhibitory signalling to the Arp2/3 complex steers cell migration. Nature 503, 281–284 (2013).

11 Liu, X. et al. Aberrant expression of Arpin in human breast cancer and its clinical significance. Journal of Cellular and Molecular Medicine 20, 450–458 (2016).

12 Lomakina, M. E. et al. Arpin downregulation in breast cancer is associated with poor prognosis. British journal of cancer 114, 545–553 (2016).

13 Li, T. et al. Clinicopathological and prognostic significance of aberrant Arpin expression in gastric cancer. World Journal of Gastroenterology 23, 1450 (2017).

14 Li, Y. et al. Restoration of Arpin suppresses aggressive phenotype of breast cancer cells. Biomedicine & Pharmacotherapy 92, 116–121 (2017).

15 Dang, I. et al. The Arp2/3 inhibitory protein Arpin is dispensable for chemotaxis. Biology of the Cell 109, 162–166 (2017).

16 Luan, Q. & Nolen, B. J. Structural basis for regulation of Arp2/3 complex by GMF. Nature structural & molecular biology 20, 1062–1068 (2013).

17 Ydenberg, C. A. et al. GMF severs actin-Arp2/3 complex branch junctions by a cofilin-like mechanism. Current biology 23, 1037–1045 (2013).

18 Lane, J., Martin, T., Weeks, H. P. & Jiang, W. G. Structure and role of WASP and WAVE in Rho GTPase signalling in cancer. Cancer genomics & proteomics 11, 155–165 (2014).

19 Fetics, S. et al. Hybrid structural analysis of the Arp2/3 regulator arpin identifies its acidic tail as a primary binding epitope. Structure 24, 252–260 (2016).

20 Zimmet, A. et al. Cryo-EM structure of NPF-bound human Arp2/3 complex and activation mechanism. Science Advances 6, eaaz7651 (2020).

21 Saks, A. J., Barrie, K. R., Rebowski, G. & Dominguez, R. NPF binding to Arp2 is allosterically linked to the release of ArpC5’s N-terminal tail and conformational changes in Arp2/3 complex. Proceedings of the National Academy of Sciences 122, e2421557122 (2025).

22 Sokolova, O. S. et al. Structural basis of Arp2/3 complex inhibition by GMF, coronin, and arpin. Journal of molecular biology 429, 237–248 (2017).

23 Fregoso, F. E. et al. Molecular mechanism of Arp2/3 complex inhibition by Arpin. Nature communications 13, 628 (2022).

24 Ti, S.-C., Jurgenson, C. T., Nolen, B. J. & Pollard, T. D. Structural and biochemical characterization of two binding sites for nucleation-promoting factor WASp-VCA on Arp2/3 complex. Proceedings of the National Academy of Sciences 108, E463–E471 (2011).

25 van Eeuwen, T. et al. Transition state of Arp2/3 complex activation by actin-bound dimeric nucleation-promoting factor. Proceedings of the National Academy of Sciences 120, e2306165120 (2023).

26 Holm, L. & Laakso, L. M. Dali server update. Nucleic acids research 44, W351–W355 (2016).

27 Montoya-García, A. et al. Arpin deficiency increases actomyosin contractility and vascular permeability. Elife 12, RP90692 (2024).

28 Abramson, J. et al. Accurate structure prediction of biomolecular interactions with AlphaFold 3. Nature 630, 493–500 (2024).

29 Lim, K.-H. et al. Ubiquitin-specific protease 11 functions as a tumor suppressor by modulating Mgl-1 protein to regulate cancer cell growth. Oncotarget 7, 14441 (2016).

30 Skalniak, L. et al. Small-molecule inhibitors of PD-1/PD-L1 immune checkpoint alleviate the PD-L1-induced exhaustion of T-cells. Oncotarget 8, 72167 (2017).

31 Soremekun, O. S., Olotu, F. A., Agoni, C. & Soliman, M. E. Recruiting monomer for dimer formation: resolving the antagonistic mechanisms of novel immune check point inhibitors against programmed death ligand-1 in cancer immunotherapy. Molecular Simulation 45, 777–789 (2019).

32 Rouiller, I. et al. The structural basis of actin filament branching by the Arp2/3 complex. The Journal of cell biology 180, 887–895 (2008).

33 Xu, X. P. et al. Three-dimensional reconstructions of Arp2/3 complex with bound nucleation promoting factors. The EMBO journal 31, 236–247 (2012).

34 Rodnick-Smith, M., Luan, Q., Liu, S.-L. & Nolen, B. J. Role and structural mechanism of WASP-triggered conformational changes in branched actin filament nucleation by Arp2/3 complex. Proceedings of the National Academy of Sciences 113, E3834–E3843 (2016).

35 Robinson, R. C. et al. Crystal structure of Arp2/3 complex. science 294, 1679–1684 (2001).

36 Smith, B. A., Daugherty-Clarke, K., Goode, B. L. & Gelles, J. Pathway of actin filament branch formation by Arp2/3 complex revealed by single-molecule imaging. Proceedings of the National Academy of Sciences 110, 1285–1290 (2013).

37 Smith, B. A. et al. Three-color single molecule imaging shows WASP detachment from Arp2/3 complex triggers actin filament branch formation. Elife 2, e01008 (2013).

38 Jasnin, M. et al. The architecture of traveling actin waves revealed by cryo-electron tomography. Structure 27, 1211–1223. e1215 (2019).

