## Supporting materials and methods, Tables S1 to S2, Figures S1 to S12 and Movies Legends for S1 for "Homodimerized Arpins: Binding to Arp2/3 Complexes with Positive Synergy to Inhibit Lamellipodia-Dependent Tumor Metastasis"

Sheng Ye

 (S. Ye)

**This PDF file includes:**

Supporting materials and methods

Tables S1 to S2

Figures S1 to S12

Movies Legends for S1

#### Crystallographic studies on Arpin

Both N- and C-terminal truncations with various lengths were screened to avoid the interference in the crystallization process, and a construct containing residues 9-221 eventually resulted in crystals, which, however, had a poor x-ray diffraction capability and could not be further optimized. Therefore, we introduced a single site-directed mutation of C133S to avoid unnecessary surface interactions and then obtained Arpin crystals in the space group of P6_5_ that diffracted to a highest resolution of 1.65 Å. To solve the phasing problem, Arpin was first subjected to a Blastp run against the Protein Data Bank (PDB) to find a search model for the molecular replacement method. Nevertheless, no structures with high similarities could be found, except the reported C-terminal tail structure of *hs*Arpin (PDB entry: 4Z68). Then, we soaked Arpin crystals in heavy-atom solutions and successfully introduced mercury ions into the crystals. Eventually, a single-wavelength dispersion dataset of a Hg derivative Arpin crystal was collected and processed to 2.1 Å. The anomalous scattering signals were strong enough to locate the Hg sites but insufficient to solve the phases. Therefore, the native dataset was combined with the Hg derivative one to perform single isomorphous replacement phasing, which successfully led to proper initial phases and a subsequent initial model. The model of Arpin was subsequently refined against the 1.65-Å native dataset to a final *R*_work_/*R*_free_ of 0.167/0.196. Crystallographic statistics are given in Table 1.

#### Multiple sequence alignment of Arpin

According to the multiple sequence alignment of Arpins from different species, there are three strictly conserved motifs (referred to as Motif 1 to 3, respectively). Both Motif 1 and 2 belong to a segment between β-strands β3 and β8 (residues 72-137) that actually includes most of the appendages to the β-barrel (helices α2, α3 and η2 and β-strands β4, β5, β6 and β7). Motif 1 (^77^KGNEIEPNFS^86^) is implicated in the formation of a β-hairpin β4-β5, which is the most flexible region of Arpin and adopts different conformations in both protomers. Motif 2 (^90^KVNTGFLMSSYKVEAKG^106^) contains β6, η2 and β7, which form a hairpin structure folded back to the surface of the central β-barrel. The two β-strands, β6 and β7, further compose a 3-stranded β-sheet with the C-terminal β-strand β11 (β6/7/11 hereafter), such that the C-terminus is tightly clamped between the β-barrel and the folded hairpin structure. In addition, a disulfide bond is formed between Cys110 and Cys183, which might not exist in cytosolic Arpins and is not conserved among species. It is interesting to note that β11 is located in Motif 3 (^178^GTVTKCNF^185^), making β-sheet β6/7/11 the most conserved region of Arpin. In fact, β6/7/11 are deeply involved in the homodimerization of Arpin by forming the “fin keel” structure, indicating an important role for Motif 2 and 3.

#### Structural conservation of Arpin

Based on the structure of Arpin, we succeeded to identify a number of unique features, including two extra regions as appendix to the central OB-fold, and the previously unknown dimeric state of Arpin with a novel homodimerization pattern. These facts urged us to find out whether the structural features we observed can be shared by Arpins from human and other species.

To assess the structural conservation of Arpin, we submitted the structure of Arpin to ConSurf server. Searching against a non-redundant protein sequence database led to 143 unique hits, including Arpins from different species, as well as predicted Arpins. Based on multiple sequence alignment of the hits, conservation scores of each residue were subsequently calculated via Bayesin method^1^, resulting in values ranging from -1.2 to 2.9, where negative and positive values represent conserved and variable residues, respectively. Since the scores provided by ConSurf have been normalized with an average value of 0 and a standard deviation of 1, the result range indicates that the sequence of Arpin have more conserved residues than variable ones.

As expected, the β-barrel core of OB-fold is relatively conserved. At the homodimeric interface, contact clusters I and II exhibit distinguished conservation degrees. Residues around contact cluster I, as well as the residues constituting the hydrophilic groove below it, are mostly variable among species, even harboring the highest conservation score in Arpin residues (2.9 for Arg135). In contrast, residues around contact cluster II are generally conserved, especially around the hydrophobic pocket and the protruding extra β-sheet β6/7/11, indicating that the extensive interprotomer interactions around cluster II are also conserved among species. Since, comparing to cluster I, contact cluster II contributes more to the formation of Arpin homodimers, we reasoned that the homodimeric form of Arpin is highly possible to be shared by Arpins from human and other species.

#### Transfection of cell lines

Transient transfection of the human fibrosarcoma HT1080 cells was performed using lipofectamine 2000 (Invitrogen, USA) according to the manufacturer’s protocol. Briefly, 1 μg plasmid DNA encoding mScarlet-tagged Arpin variants (pcDNA3.1 vector) was diluted in 150 μL Opti-MEM to prepare the plasmid solution. 3 μL Lipofectamine 2000 was diluted in 150 μL Opti-MEM to prepare the liposome solution. Both mixtures were incubated at room temperature for 5 min. The plasmid solution was then added dropwise to the liposome solution and incubated for 20 min at room temperature. The resulting transfection mixture was added dropwise to 10 cm dishes containing HT1080 cells at approximately 80% confluency. Six hours post-transfection, the culture medium was replaced with high-glucose DMEM lacking glutamine and supplemented with 10% FBS. Protein expression was assessed 24 h later by flow cytometry.

HT1080 cells with stable mScarlet-tagged Arpin variants expression were also generated by lentiviral infection following the appropriate biological safety regulations. To deliver plasmids to generate the lentiviral particles encoding protein, lipofectamine 2000 was used according to the manufacturer´s protocol. 3.5μg of the plasmid pCDH-Arpin variants encoding the mScarlet protein was incubated with packaging plasmids (2.5 μg pLP1, 2.5 μg pLP2, 1.5 μg pLP-VSVG) in 200 μl Opti-MEM. The mixture was vortexed for 10 s. In a separate mix, 15 μl of lipofectamine 2000 was added to 200 μl Opti-MEM and was gently added to the first mixture followed by 5 min incubation at room temperature. The transfection mix was added drop-wise to a 10 cm dish of ~80% confluent HT1080 cells. 6 h post-transfection, the medium was replaced with high-glucose DMEM without glutamine, supplemented with 10% FBS. 48 h later, the lentiviral medium was collected and filtered through 0.45 µm filters. Target cells were spinfected with the optimized titer of the lentiviral medium by centrifugation at 4000 × rpm for 10 min at room temperature. Spinfected cells were recovered for one day in fresh medium and mScarlet expression was confirmed by flow cytometry. Lentiviral transduction was performed by adding viral particles to cells in culture medium at a 1:1 ratio (virus: DMEM) supplemented with polybrene (final concentration: 8 μg/mL). After 12 h incubation, the medium was replaced with fresh DMEM containing 10% FBS. Expression of the target gene with fluorescent tag was assessed 48 h infection. Stable cell lines were generated by subsequent puromycin selection. Cells were initially treated with 2 μg/mL puromycin, with the concentration gradually increased (up to 15 μg/mL) over several passages until a fully positive, stably expressing population was obtained.

#### SIM Imaging of MCF7 Cells

SIM images of MCF7 cells were acquired on the DeltaVision OMX V3 imaging system (GE Healthcare) with a ×100/1.40 NA oil objective (Olympus UPlanSApo), solid-state multimode lasers (488, 561nm) and electron-multiplying CCD (charge-coupled device) cameras (Evolve 512×512, Photometrics). Serial Z-stack sectioning was done at 125 nm intervals for SIM mode. To obtain optimal images, immersion oils with refractive indices of 1.516 were used for MCF7 cells on glass coverslips. The microscope is routinely calibrated with 100 nm fluorescent spheres to calculate both the lateral and axial limits of image resolution. SIM image stacks were reconstructed using softWoRx 6.1.1 (GE Healthcare) with the following settings: pixel size 39.5 nm; channel-specific optical transfer functions; Wiener filter constant 0.0010; discard Negative Intensities background; drift correction with respect to first angle; custom K0 guess angles for camera positions. The reconstructed images were further processed for maximum-intensity projections with softWoRx 6.1.1. Pixel registration was corrected to be less than 1 pixel for all channels using 100 nm Tetraspeck beads.

#### Live cell time-lapse imaging and trajectory analysis

An PE Operetta High Content Screening System (Perkin Elmer, USA) was used in wide field mode (×10 objective) to capture 9 tiles within each 96-well while simultaneously acquiring 3 channels (Bright field observation, Differential phase contrast (DPC), mScarlet). For Movie S1, the images were taken once in 10 min for 6 hours. Image analysis was performed using Harmony 4.0 software, enabling automated imaging then analysis during the screening process. The main steps of the analysis protocol involved identifying transfected cells via the red fluorescence channel and quantifying total cell counts per well using the DPC channel. To minimize positional drift caused by early plate movement, imaging was initiated from the fifth time point, when the plate was stably settled on the stage. The following criteria were obtained at all time points, including accumulated distance (μm), displacement (μm), speed (μm/s), and directional persistence (straightness). In a protocol refinement step, cells were filtered by selecting those with "Begin" as the start type and "End" as the end type prior to statistical analysis. Cells labeled as Split, Border, Merged, or Cosmos during tracking were excluded to minimize artifacts and ensure accurate quantification of individual cell trajectories.

#### AI-aided design of the dual-tailed peptide

Due to the absence of high-resolution experimental structure for the Homo sapiens Arpin-bound Arp 2/3 complex, the homologous structure of Actin obtained from Bos taurus was employed as the foundational template. The cryo-EM-derived structure, PDB entry 7JPN (resolution: 3.2 Å), was selected as the initial model, which represents the assembly state of the Arpin-actin complex.

For computational design, the α-helical region (residues number 194–206, as shown in Figure S1, the canonical binding site) was selected as the region to be redesigned. For this helical segment, over 600 electron densities were produced using our proprietary generative algorithm. With iterative refinement, 2,292 structural samples were obtained through employing our in-house developed Deep-learning-based segmentation model to identify the residue type and conformation.

The CryoEM structure of the Arp2/3complex with bound NPFs (PDB entry: 6UHC, resolution: 3.9 Å) and Arpin (PDB 7JPN, resolution: 3.2 Å) was utilized as a reference when there were no human Arpin-bound complex structures at both biding sites. Potential secondary binding sites for Arpin were identified by structurally superimposing the two complexes. This aligned structure, as shown in Figure S12, served as the starting point for subsequent computational screening, with the objective of identifying designed binders with better binding affinity. Biopython-based structural analysis pipelines were used to quantify hydrogen bonds, steric conflicts, salt bridge interactions, and buried solvent-accessible surface area (SASA), and absolute binding free energies were computed using the APBS (Adaptive Poisson-Boltzmann Solver) suite.

In order to prioritize high-probability binding candidates, structural evaluations were conducted on both the canonical binding site and the putative secondary binding site, incorporating both geometric and energetic factors. Finally, six candidates were selected for further experimental validation. The details with the sequence and the geometric and energetic parameters for the suspected secondary binding as well as canonical binding show in Table S1 and S2.

### References

1	Ben Chorin, A. *et al.* ConSurf‐DB: An accessible repository for the evolutionary conservation patterns of the majority of PDB proteins. *Protein Science* **29**, 258-267 (2020).

### Table S1. Computational screening results at the canonical binding site.

| ID | sequence | contacts | Salt bridges | clashes | binding_energy  （kcal/mol） |
| --- | --- | --- | --- | --- | --- |
| dP_arpin_0001 | QWKTKVANKAHQA | 83 | 0 | 0 | -10.679 |
| dP_arpin_0002 | EWKTKVANKAHQA | 83 | 0 | 0 | -21.9851 |
| dP_arpin_0003 | QWKTKVADKAHQA | 83 | 0 | 0 | -14.9045 |
| dP_arpin_0004 | EWKTKVADKAHQA | 83 | 0 | 0 | -24.3996 |
| dP_arpin_0005 | QWKTKVANKAHEA | 83 | 0 | 0 | -15.4532 |
| dP_arpin_0006 | EWKTKVANKAHEA | 83 | 0 | 0 | -26.3723 |
| dP_arpin_0007 | QWKTKVADKAHEA | 83 | 0 | 0 | -18.5115 |
| dP_arpin_0008 | EWKTKVADKAHEA | 83 | 0 | 0 | -27.6783 |
| dP_arpin_0009 | FWKSVVASQKQQQ | 138 | 0 | 0 | -380.142 |
| dP_arpin_0010 | FWKSVVASEKQQQ | 138 | 0 | 0 | -216.377 |
| dP_arpin_0011 | FWKSVVASQKEQQ | 138 | 0 | 0 | -375.764 |
| dP_arpin_0012 | FWKSVVASEKEQQ | 138 | 0 | 0 | -211.026 |
| dP_arpin_0013 | FWKSVVASQKQEQ | 138 | 0 | 0 | -77.7391 |
| dP_arpin_0014 | FWKSVVASEKQEQ | 138 | 0 | 0 | -196.761 |
| dP_arpin_0015 | FWKSVVASQKEEQ | 138 | 0 | 0 | -73.1259 |
| dP_arpin_0016 | FWKSVVASEKEEQ | 138 | 0 | 0 | -191.381 |
| dP_arpin_0017 | AVIQQVNKASQQA | 39 | 0 | 0 | 180.4249 |
| dP_arpin_0018 | AVIEQVNKASQQA | 39 | 0 | 0 | 168.2103 |
| dP_arpin_0019 | AVIQEVNKASEQA | 39 | 0 | 0 | 202.8576 |
| dP_arpin_0020 | AVIEEVNKASEQA | 39 | 0 | 0 | 192.5639 |
| dP_arpin_0021 | AVIQQVDKASEQA | 39 | 2 | 0 | 185.2027 |
| dP_arpin_0022 | AVIEQVDKASEQA | 39 | 2 | 0 | 173.8152 |
| dP_arpin_0023 | AVIQEVDKASEQA | 39 | 2 | 0 | 207.0622 |
| dP_arpin_0024 | AVIEEVDKASEQA | 39 | 2 | 0 | 197.3668 |
| dP_arpin_0025 | AVIQEVNKASQQA | 39 | 0 | 0 | 200.294 |
| dP_arpin_0026 | AVIEEVNKASQQA | 39 | 0 | 0 | 189.9707 |
| dP_arpin_0027 | AVIQQVDKASQQA | 39 | 2 | 0 | 182.3052 |
| dP_arpin_0028 | AVIEQVDKASQQA | 39 | 2 | 0 | 170.8352 |
| dP_arpin_0029 | AVIQEVDKASQQA | 39 | 2 | 0 | 203.8147 |
| dP_arpin_0030 | AVIEEVDKASQQA | 39 | 2 | 0 | 194.1416 |
| dP_arpin_0031 | AVIQQVNKASEQA | 39 | 0 | 0 | 182.5513 |
| dP_arpin_0032 | AVIEQVNKASEQA | 39 | 0 | 0 | 170.457 |
| dP_arpin_0033 | SLIAKVQKLVVAK | 71 | 0 | 3 | 1.535628 |
| dP_arpin_0034 | SLIAKVEKLVVAK | 71 | 2 | 3 | 0.814553 |
| dP_arpin_0535 | ATKKVATAQYQNS | 61 | 0 | 0 | 57.52925 |
| dP_arpin_0536 | ATKKVATAQYENS | 61 | 0 | 0 | 59.23949 |
| dP_arpin_0537 | ATKKVATAQYQDS | 61 | 0 | 0 | 60.77753 |
| dP_arpin_0538 | ATKKVATAQYEDS | 61 | 0 | 0 | 62.51651 |
| dP_arpin_0539 | KLVQTVTSLKTRS | 51 | 0 | 0 | 195.9413 |
| dP_arpin_0540 | KLVETVTSLKTRS | 51 | 0 | 0 | 197.7891 |
| dP_arpin_0541 | KVNHQVVKIHAAS | 55 | 0 | 0 | 459.8654 |
| dP_arpin_0542 | KVDHQVVKIHAAS | 55 | 0 | 0 | 489.6458 |
| dP_arpin_0543 | KVNHEVVKIHAAS | 55 | 0 | 0 | 468.3837 |
| dP_arpin_0544 | KVDHEVVKIHAAS | 55 | 0 | 0 | 499.7697 |
| dP_arpin_0545 | SLVVKVSAASNRA | 47 | 0 | 0 | 202.571 |
| dP_arpin_0546 | SLVVKVSAASDRA | 47 | 0 | 0 | 202.0906 |
| dP_arpin_0547 | NLKKIVAAIASFA | 50 | 0 | 0 | 163.7491 |
| dP_arpin_0548 | DLKKIVAAIASFA | 50 | 0 | 0 | 81.84044 |
| dP_arpin_0549 | DLCIKTANLAEAV | 49 | 0 | 0 | 205.747 |
| dP_arpin_0550 | KLSKTSAALAVSA | 39 | 0 | 0 | 244.2823 |
| dP_arpin_0551 | QLVASVATLSATS | 47 | 0 | 0 | 295.1066 |
| dP_arpin_0552 | ELVASVATLSATS | 47 | 0 | 0 | 295.5443 |
| dP_arpin_0553 | SLKAKVSKQKALK | 58 | 0 | 0 | 283.6236 |
| dP_arpin_0554 | SLKAKVSKEKALK | 58 | 0 | 0 | 292.9358 |
| dP_arpin_1055 | DVVRKVNQLAKSA | 41 | 0 | 0 | 88.29351 |
| dP_arpin_1056 | DVVRKVDQLAKSA | 41 | 2 | 0 | 96.35349 |
| dP_arpin_1057 | DVVRKVNELAKSA | 41 | 0 | 0 | 90.18701 |
| dP_arpin_1058 | DVVRKVDELAKSA | 41 | 2 | 0 | 99.40566 |
| dP_arpin_1059 | QIIAVVALTAVLV | 49 | 0 | 0 | 68.63249 |
| dP_arpin_1060 | EIIAVVALTAVLV | 49 | 0 | 0 | 66.52035 |
| dP_arpin_1061 | NVKALSAKKVFAS | 52 | 0 | 0 | 138.4554 |
| dP_arpin_1062 | DVKALSAKKVFAS | 52 | 0 | 0 | 184.0313 |
| dP_arpin_1063 | NVVRKVNQLAKSA | 41 | 0 | 0 | 107.4244 |
| dP_arpin_1064 | DVVRKVNQLAKSA | 41 | 0 | 0 | 88.29351 |
| dP_arpin_1065 | NVVRKVDQLAKSA | 41 | 2 | 0 | 114.7842 |
| dP_arpin_1066 | DVVRKVDQLAKSA | 41 | 2 | 0 | 96.35349 |
| dP_arpin_1067 | NVVRKVNELAKSA | 41 | 0 | 0 | 108.4894 |
| dP_arpin_1068 | DVVRKVNELAKSA | 41 | 0 | 0 | 90.18701 |
| dP_arpin_1069 | NVVRKVDELAKSA | 41 | 2 | 0 | 117.022 |
| dP_arpin_1070 | DVVRKVDELAKSA | 41 | 2 | 0 | 99.40566 |
| dP_arpin_1071 | AVVVVVARIVKNS | 37 | 0 | 0 | 190.5833 |
| dP_arpin_1072 | AVVVVVARIVKDS | 37 | 0 | 0 | 196.3851 |
| dP_arpin_1073 | DQVVAVAKVQYAA | 44 | 0 | 0 | 167.7675 |
| dP_arpin_1074 | NQVVAVAKVEYAA | 44 | 0 | 0 | 190.3265 |
| dP_arpin_1075 | DQVVAVAKVEYAA | 44 | 0 | 0 | 110.1761 |
| dP_arpin_1076 | DEVVAVAKVEYAA | 44 | 0 | 0 | 186.7769 |
| dP_arpin_1077 | AVVIVVASFIKQS | 37 | 0 | 0 | 111.8765 |
| dP_arpin_1078 | QVIQVVAVNVSAA | 41 | 0 | 0 | 139.9893 |
| dP_arpin_1079 | EVIQVVAVNVSAA | 41 | 0 | 0 | 137.2367 |
| dP_arpin_1080 | QVIEVVAVNVSAA | 41 | 0 | 0 | 142.2389 |
| dP_arpin_1081 | EVIEVVAVNVSAA | 41 | 0 | 0 | 140.2551 |
| dP_arpin_1082 | AVIIAVASIKARV | 55 | 0 | 0 | 198.9389 |

### Table S2. Computational screening results at the putative secondary binding site.

| ID | contacts | saltbridges | clashes | buried_SA | binding_energy  （kcal/mol） |
| --- | --- | --- | --- | --- | --- |
| dP_arpin_0001 | 37 | 0 | 1 | 10.35791 | 183.9926506 |
| dP_arpin_0002 | 37 | 0 | 1 | 10.35791 | 182.9153402 |
| dP_arpin_0003 | 37 | 0 | 1 | 10.35785 | 187.252262 |
| dP_arpin_0004 | 37 | 0 | 1 | 10.35785 | 186.4054686 |
| dP_arpin_0005 | 37 | 0 | 1 | 10.35492 | 199.9158263 |
| dP_arpin_0006 | 37 | 0 | 1 | 10.35492 | 199.1863452 |
| dP_arpin_0007 | 37 | 0 | 1 | 10.35498 | 204.4795064 |
| dP_arpin_0008 | 37 | 0 | 1 | 10.35498 | 203.8809939 |
| dP_arpin_0009 | 75 | 1 | 4 | 10.99414 | 90.38109819 |
| dP_arpin_0010 | 75 | 1 | 4 | 10.9903 | 104.7014706 |
| dP_arpin_0011 | 75 | 1 | 4 | 10.99414 | 87.94703737 |
| dP_arpin_0012 | 75 | 1 | 4 | 10.9903 | 102.4290464 |
| dP_arpin_0013 | 75 | 1 | 4 | 10.98547 | 120.273144 |
| dP_arpin_0014 | 75 | 1 | 4 | 10.98169 | 135.7710195 |
| dP_arpin_0015 | 75 | 1 | 4 | 10.98547 | 118.7357 |
| dP_arpin_0016 | 75 | 1 | 4 | 10.98169 | 134.1918801 |
| dP_arpin_0017 | 30 | 0 | 1 | 9.324158 | -81.64570311 |
| dP_arpin_0018 | 30 | 0 | 1 | 9.324097 | -81.14807469 |
| dP_arpin_0019 | 30 | 0 | 1 | 9.303528 | -68.92425089 |
| dP_arpin_0020 | 30 | 0 | 1 | 9.303589 | -68.27339121 |
| dP_arpin_0021 | 30 | 0 | 1 | 9.303528 | -67.0738553 |
| dP_arpin_0022 | 30 | 0 | 1 | 9.303528 | -66.49799157 |
| dP_arpin_0023 | 30 | 0 | 1 | 9.303589 | -64.94197463 |
| dP_arpin_0024 | 30 | 0 | 1 | 9.303589 | -64.38404841 |
| dP_arpin_0025 | 30 | 0 | 1 | 9.324097 | -80.14459939 |
| dP_arpin_0026 | 30 | 0 | 1 | 9.324097 | -79.52570526 |
| dP_arpin_0027 | 30 | 0 | 1 | 9.324097 | -78.10828385 |
| dP_arpin_0028 | 30 | 0 | 1 | 9.324097 | -77.55635139 |
| dP_arpin_0029 | 30 | 0 | 1 | 9.324097 | -76.41449659 |
| dP_arpin_0030 | 30 | 0 | 1 | 9.324097 | -75.80491422 |
| dP_arpin_0031 | 30 | 0 | 1 | 9.303589 | -70.9444145 |
| dP_arpin_0032 | 30 | 0 | 1 | 9.303589 | -70.33823926 |
| dP_arpin_0033 | 25 | 0 | 1 | 8.028564 | 122.5227256 |
| dP_arpin_0034 | 25 | 0 | 1 | 8.028564 | 123.2858531 |
| dP_arpin_0535 | 25 | 1 | 0 | 8.756409 | 167.7423626 |
| dP_arpin_0536 | 25 | 1 | 0 | 8.741455 | 164.1490145 |
| dP_arpin_0537 | 25 | 1 | 0 | 8.750732 | 180.6458869 |
| dP_arpin_0538 | 25 | 1 | 0 | 8.735718 | 179.2961496 |
| dP_arpin_0539 | 42 | 0 | 2 | 9.166687 | 98.03600225 |
| dP_arpin_0540 | 42 | 0 | 2 | 9.166687 | 99.16010505 |
| dP_arpin_0541 | 15 | 1 | 0 | 8.383606 | -580.0564499 |
| dP_arpin_0542 | 15 | 1 | 0 | 8.379578 | -570.0738834 |
| dP_arpin_0543 | 15 | 1 | 0 | 8.381042 | -574.5371752 |
| dP_arpin_0544 | 15 | 1 | 0 | 8.376953 | -564.4343432 |
| dP_arpin_0545 | 13 | 0 | 0 | 7.851135 | 85.84630573 |
| dP_arpin_0546 | 13 | 0 | 0 | 7.851135 | 84.48292517 |
| dP_arpin_0547 | 31 | 1 | 2 | 8.798584 | 23.67949181 |
| dP_arpin_0548 | 31 | 1 | 2 | 8.798523 | 21.27153368 |
| dP_arpin_0549 | 12 | 0 | 0 | 8.469849 | 99.03573351 |
| dP_arpin_0550 | 10 | 0 | 0 | 7.156982 | -79.79173256 |
| dP_arpin_0551 | 11 | 0 | 0 | 8.224426 | 68.87982936 |
| dP_arpin_0552 | 11 | 0 | 0 | 8.224426 | 67.72872998 |
| dP_arpin_0553 | 26 | 2 | 0 | 10.3382 | -218.2488409 |
| dP_arpin_0554 | 26 | 2 | 0 | 10.33588 | -213.8005377 |
| dP_arpin_1055 | 8 | 0 | 0 | 7.131409 | 82.17364736 |
| dP_arpin_1056 | 8 | 0 | 0 | 7.13147 | 86.35067584 |
| dP_arpin_1057 | 8 | 0 | 0 | 7.131409 | 86.44700275 |
| dP_arpin_1058 | 8 | 0 | 0 | 7.131409 | 90.95290242 |
| dP_arpin_1059 | 20 | 0 | 0 | 8.378906 | 133.9748762 |
| dP_arpin_1060 | 20 | 0 | 0 | 8.378967 | 132.1958788 |
| dP_arpin_1061 | 12 | 0 | 0 | 9.645691 | 81.19163849 |
| dP_arpin_1062 | 12 | 0 | 0 | 9.645691 | 78.96280906 |
| dP_arpin_1063 | 8 | 0 | 0 | 7.131409 | 78.80420451 |
| dP_arpin_1064 | 8 | 0 | 0 | 7.131409 | 82.17364736 |
| dP_arpin_1065 | 8 | 0 | 0 | 7.13147 | 82.94504654 |
| dP_arpin_1066 | 8 | 0 | 0 | 7.13147 | 86.35067584 |
| dP_arpin_1067 | 8 | 0 | 0 | 7.131409 | 82.87029559 |
| dP_arpin_1068 | 8 | 0 | 0 | 7.131409 | 86.44700275 |
| dP_arpin_1069 | 8 | 0 | 0 | 7.131409 | 87.37782423 |
| dP_arpin_1070 | 8 | 0 | 0 | 7.131409 | 90.95290242 |
| dP_arpin_1071 | 26 | 4 | 0 | 9.43042 | -80.00241964 |
| dP_arpin_1072 | 26 | 4 | 0 | 9.419434 | -59.73026003 |
| dP_arpin_1073 | 19 | 0 | 0 | 7.961731 | 71.94127255 |
| dP_arpin_1074 | 19 | 0 | 0 | 7.957153 | 77.40612268 |
| dP_arpin_1075 | 19 | 0 | 0 | 7.957214 | 75.86595646 |
| dP_arpin_1076 | 19 | 2 | 0 | 7.949463 | 66.15939193 |
| dP_arpin_1077 | 33 | 0 | 2 | 9.453308 | 39.88748269 |
| dP_arpin_1078 | 5 | 0 | 0 | 7.743713 | 93.99354968 |
| dP_arpin_1079 | 5 | 0 | 0 | 7.743774 | 95.00732732 |
| dP_arpin_1080 | 5 | 0 | 0 | 7.743774 | 95.45144708 |
| dP_arpin_1081 | 5 | 0 | 0 | 7.743713 | 96.4544793 |
| dP_arpin_1082 | 37 | 2 | 1 | 9.44574 | -211.3670536 |

### Figure Legends

**Figure S1. Secondary structure and sequence alignment of Arpin from different species**. Including *Danio rerio* (NM_001017780), *Xenopus tropicalis* (NM_001004898), *Bos taurus* (NM_001076304), *Mus musculus* (NM_027420), *Homo sapiens* (NM_182616). Conserved amino acid sequences crossed Arpin family are indicated by red and similar residues are indicated by yellow.

**Figure S2. (A) Cartoon representation and (B) topology diagram of Arpin promoter structure.** The six helices (α1, α2, η1, η2, η3 and η4) and β-strands (β1-β11) are highlighted in ribbon (red and yellow), and the rest residues are shown in thin lines (green).

**Figure S3. Structural superposition and comparison between Arpin and similar proteins.** (A). Structural alignment between chain A and chain B in Arpin crystal structure. Arpin was compared with (B) Rim1 (PDB entry: 6CQK) and CusF (PDB entry: 3E6Z) conformations. The r.m.s. deviations are 1.95-Å for 71 aligned Cα atoms of Rim1 (z-score 6.4), and 2.41 Å for 68 aligned Cα atoms of CusF (z-score 6.4).

**Figure S4.** **Stereo diagram of protein contact shows a close-up view of Cluster I area at the dimer interface.** Contact Cluster I lies between the β-barrels, where the N-terminus of one protomer interacts with the η4 helix of the other via hydrophobic contacts and hydrogen bonds.

**Figure S5. Opposite view of the conserved surface region on the dimer of Arpin.** 180° rotated surface view of the Arpin dimer highlights the conserved “fin keel” structure from the opposite side, confirming its bilateral conservation pattern.

**Figure S6.** **Enclosed interfacial cavity of the Arpin dimer.** The experimentally phased electron density reveals a sealed cavity situated between the two contact clusters. This cavity becomes isolated from bulk solvent upon dimer formation.

**Figure S7. Sensorgram summary from SPR analysis showing interactions between Arpin variants and Arp2/3 complex.** Arp2/3 complex was immobilized on CM5 surface vs increasing concentrations of Arpin variants. Biacore evaluation software 3.0 was used to fit binding model.

**Figure S8. Representative images of crystal violet staining after cell migration captured 24 hours after treatment.** Overexpression of Arpin and truncated peptide Arpin_CT_ and Arpin_AT_ (contain putative interaction domains with the Arp2/3 complex) induced inhibition of cell migration in MCF7 cells. However, the effect of this inhibitory binding was reduced when Arpin was converted from a homodimer to two protomers (Arpin^FM^). When Arpin^∆CT^ was overexpressed, the cell migrating ability rose sharply dramatically to a level comparable to that of cells treated only with pcDNA3.1.

**Figure S9. Structural comparison of Arpin globular core and GST homodimers.** (A). Structural measurement of the distance between C-terminal Cα atoms of the homodimer of Arpin globular core, approximately 25.0 Å, with both termini symmetrically arranged and oriented outward. (B). Structural measurement of the GST homodimer (PDB ID: 6JI6), showing an inter-terminal Cα distance of approximately 46.7 Å. The two C-termini also exhibit twofold rotational symmetry and extend outward, providing a favorable geometry for fusion.

**Figure S10. Covalent crosslinking of D-aCT peptides.** Schematic diagram of D-aCT construct, incorporating N-terminal cysteine residues and a flexible spacer for crosslinking.

**Figure S11. Binding affinity of D-aCT and AI-designed dual-tailed peptide to the Arp2/3 complex.** SPR analysis was performed with immobilized Arp2/3 complex on a CM5 sensor chip and increasing concentrations of dual-tailed peptide. Binding curves were fitted using Biacore Evaluation Software 3.0 to determine dissociation constants.

**Figure S12. Potential secondary binding sites of Arpin were determined by structural superposition.** The CryoEM structure of the Arp2/3complex with bound NPFs (PDB 6UHC, resolution: 3.9 Å) and Arpin (PDB 7JPN, resolution: 3.2 Å) were utilized as a reference and identified by structurally superimposing the two complexes.

**Figures**

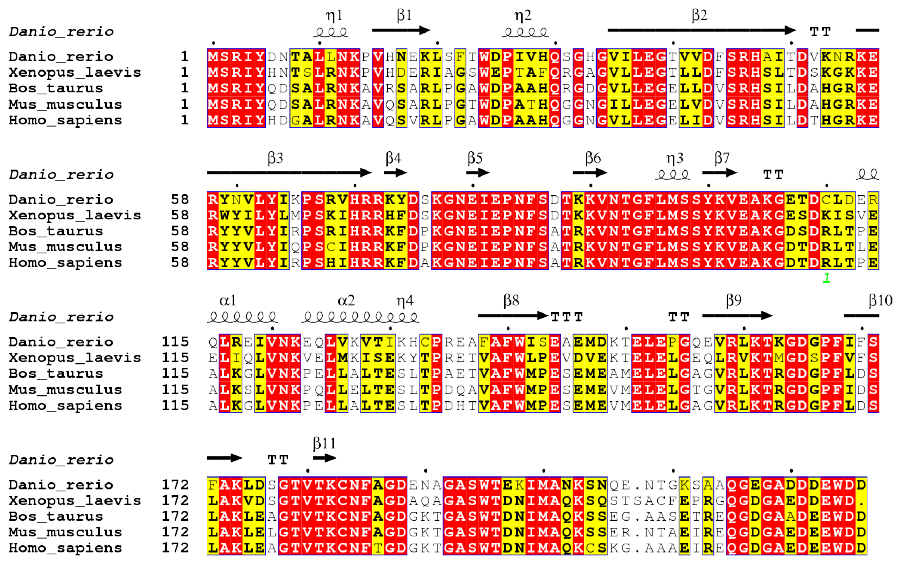

**Figure S1**

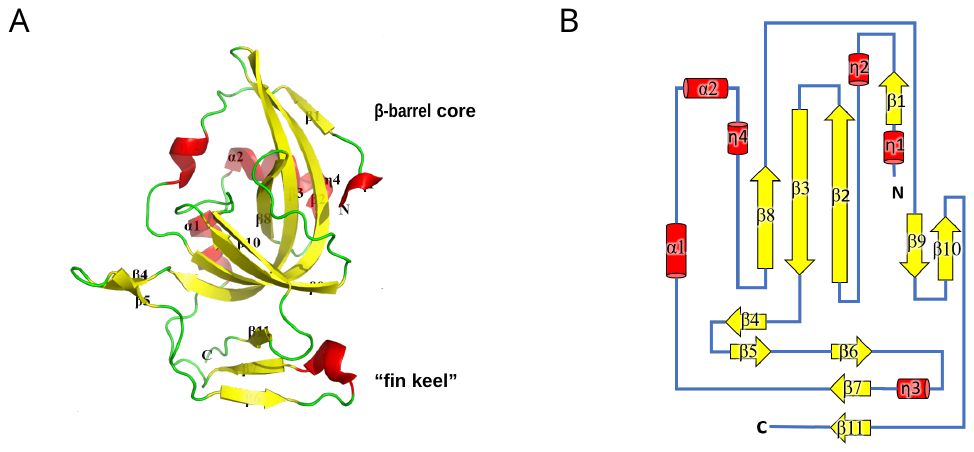

**Figure S2**

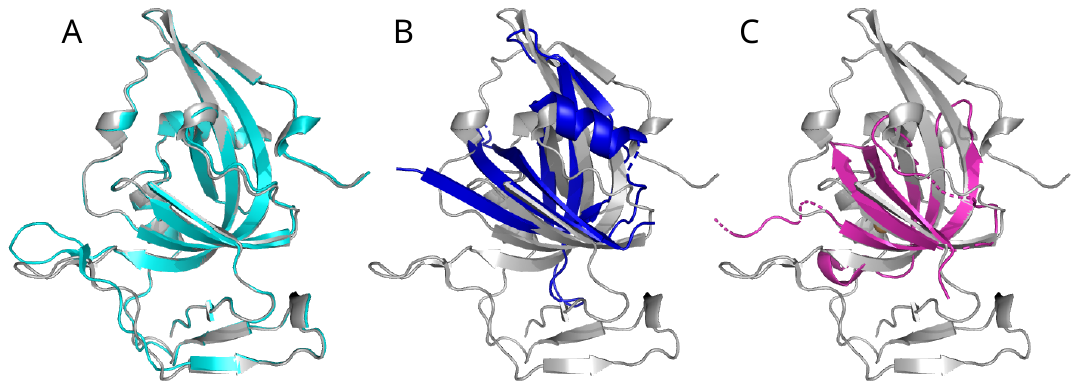

**Figure S3**

**Figure S4**

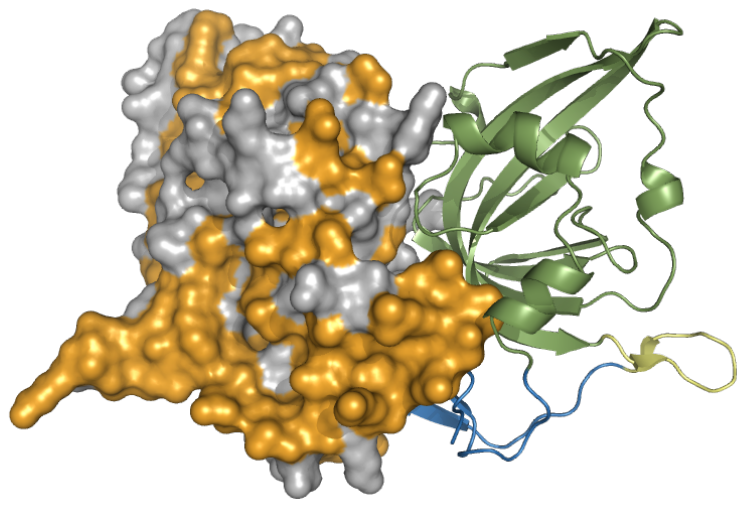

**Figure S5**

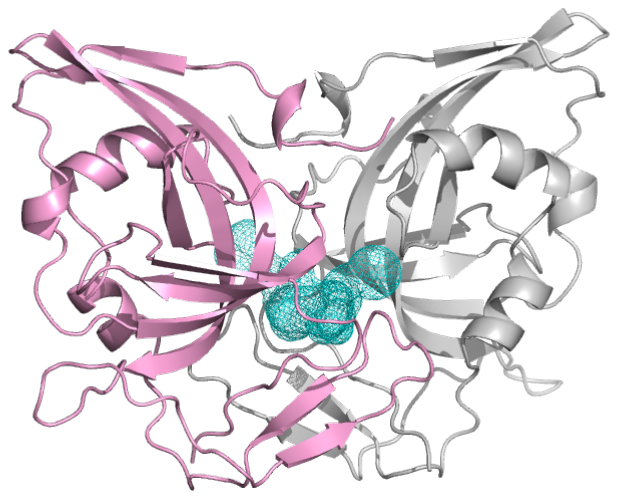

**Figure S6**

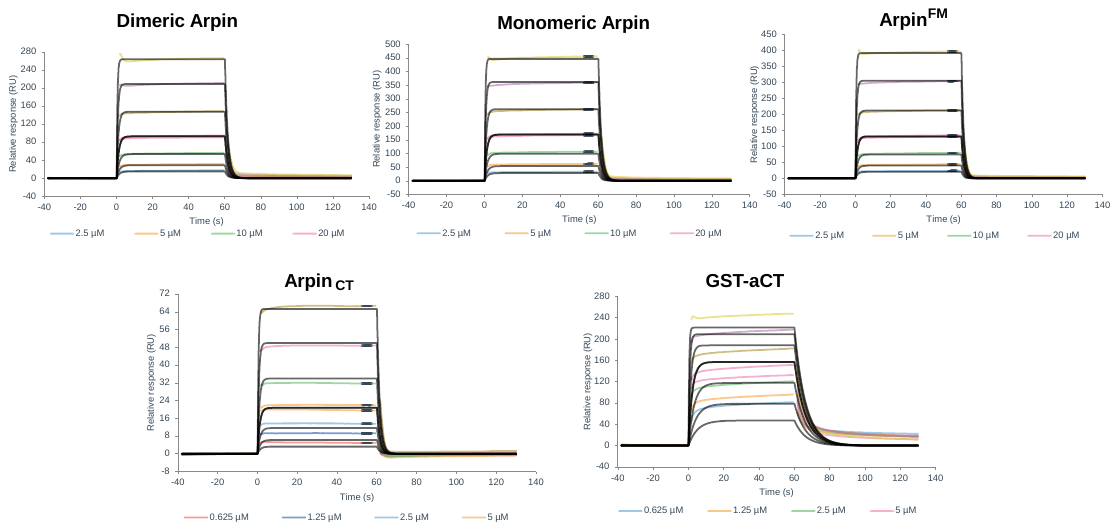

**Figure S7**

**Figure S8**

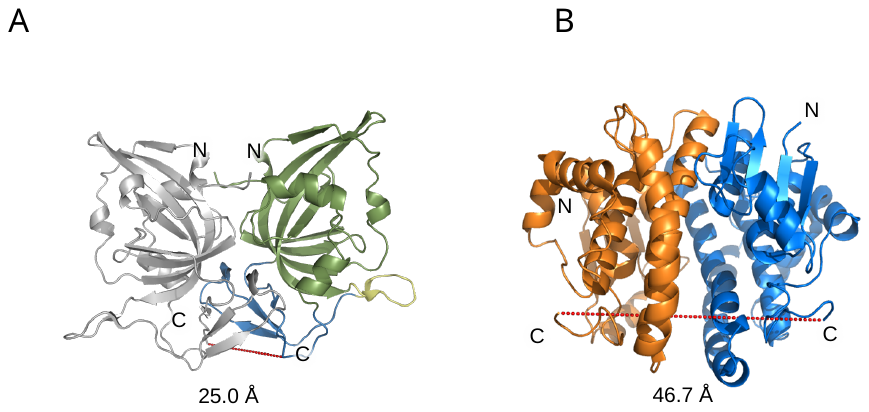

**Figure S9**

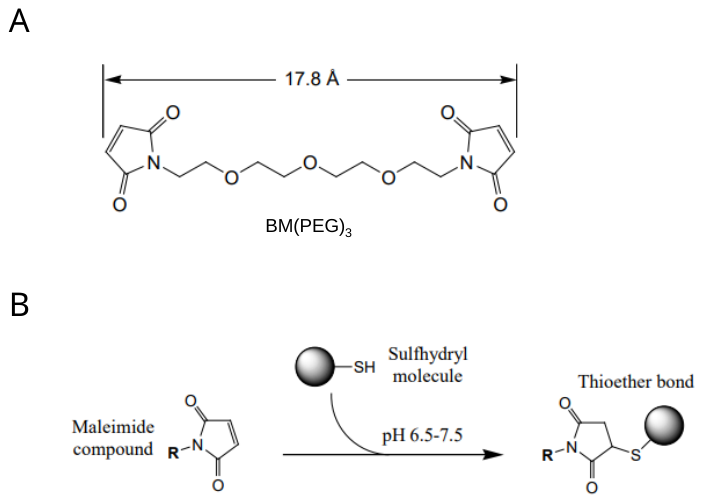

**Figure S10**

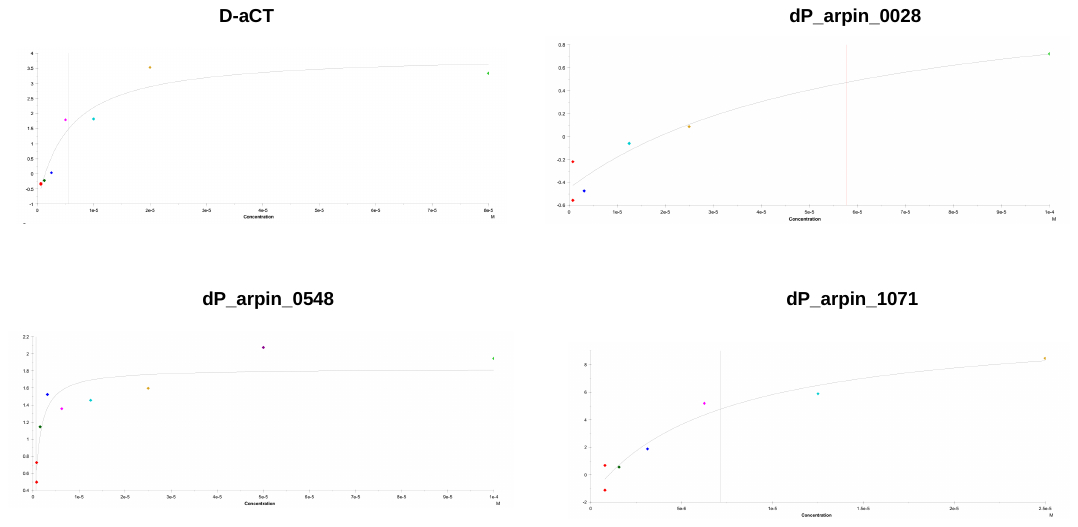

**Figure S11**

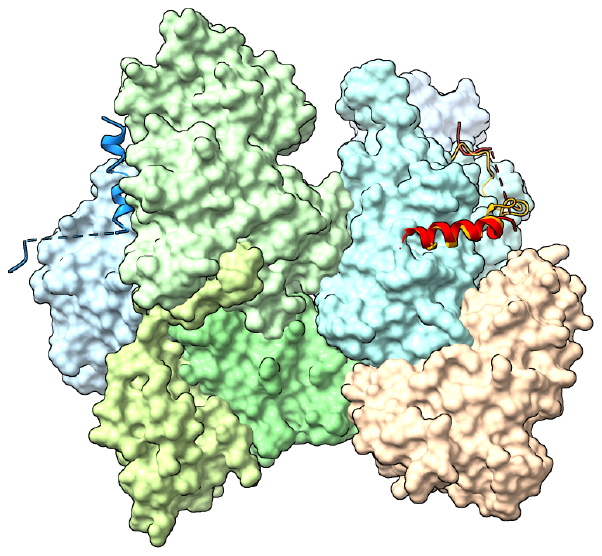

**Figure S12**

### Movie S1. Time-lapse imaging of HT1080 cell migration.

Representative 6h trajectories of single-cell migration under various Arpin construct conditions. Overexpression of full-length Arpin or engineered variants alters cell movement patterns and reduces directional persistence.
